# Cavin3 released from caveolae interacts with BRCA1 to regulate the cellular stress response

**DOI:** 10.1101/2020.07.26.222158

**Authors:** Kerrie-Ann McMahon, David A. Stroud, Yann Gambin, Vikas A. Tillu, Michele Bastiani, Emma Sierecki, Mark Polinkovsky, Thomas E. Hall, Guillermo A. Gomez, Yeping Wu, Marie-Odile Parat, Nick Martel, Harriet P. Lo, Kum Kum Khanna, Kirill Alexandrov, Roger Daly, Alpha S. Yap, Michael T. Ryan, Robert G. Parton

## Abstract

Caveolae-associated protein 3 (cavin3), a putative tumor suppressor protein, is inactivated in most cancers. We characterized how cavin3 affects the cellular proteome using genome-edited cells together with label-free quantitative proteomics. These studies revealed a prominent role for cavin3 in DNA repair with BRCA1 and BRCA1 A-complex components being downregulated on cavin3 deletion. Cellular and cell-free expression assays, we show a direct interaction between BRCA1 and cavin3. Association of BRCA1 and cavin3 occurs when cavin3 is released from caveolae that are disassembled in response to UV and mechanical stress. Supporting a role in DNA repair, cavin3-deficient cells were sensitized to the effects of PARP inhibition, which compromises DNA repair, and showed reduced recruitment of the BRCA1 A-complex to UV DNA damage foci. Overexpression and RNAi-depletion revealed that cavin3 sensitized various cancer cells to UV-induced apoptosis. We conclude that cavin3 functions together with BRCA1 in multiple pathways that contribute to tumorigenesis.

## Introduction

Caveolae are an abundant surface feature of most vertebrate cells. Morphologically, caveolae are 50-100 nm bulb-shaped structures attached to the plasma membrane (Parton and del Pozo, 2013). One of the defining features of this domain is the integral membrane protein caveolin-1 (CAV1). CAV1 is a structural component of caveolae regulating diverse cellular processes, including endocytosis, vesicular transport, cell migration, and signal transduction (Parton and del Pozo, 2013).

Recently, we and others have characterized a caveolar adaptor molecule, caveolae-associated protein 3 (cavin3) (McMahon et al., 2009). Cavin3 belongs to a family of proteins that includes caveolae-associated protein 1 (cavin1), caveolae-associated protein 2 (cavin2), and the muscle-specific member caveolae-associated protein 4 (cavin4) (Ariotti and Parton, 2013; Bastiani et al., 2009; Hansen et al., 2009, 2013, Kovtun et al., 2015; Lo et al., 2015; McMahon et al., 2009). Cavin3 is a putative tumor suppressor gene that is epigenetically silenced in a range of human malignancies (Xu et al., 2001), principally due to hypermethylation of its promoter region (Caren et al., 2011; Kim et al., 2014; Lee et al., 2008; Lee et al., 2011; Martinez et al., 2009; Tong et al., 2010; Zochbauer-Muller et al., 2005). Furthermore, cavin3 has previously suggested to interact with BRCA1, although no data has been formally published to support this interaction (Xu et al., 2001). Several studies have implicated Cavin3 in a broad range of cancer-related processes including proliferation, apoptosis, Warburg metabolism, as well as in cell migration and matrix metalloproteinase regulation; however, the molecular basis of its actions is poorly understood (Hernandez et al., 2013; Toufaily et al., 2014).

BReast CAncer gene 1 (BRCA1) is a significant breast cancer suppressor gene. It is one of the most frequently mutated genes in hereditary breast cancer (King et al., 2003; Miki et al., 1994; Venkitaraman et al., 2002). Also, BRCA1 levels are reduced or absent in many sporadic breast cancers due to gene silencing by promoter methylation or downregulation of the gene by other tumor suppressors or oncogenes (Mueller and Roskelley, 2003; Turner et al., 2004). BRCA1 has been implicated in a remarkable number of processes, including cell cycle checkpoint control, DNA damage repair, and transcriptional regulation (reviewed by Lord and Ashworth, 2016; Savage et al., 2015). At the molecular level, accumulated evidence suggests that BRCA1 plays an integral role in the formation of several macromolecular complexes (BRCA1 A, BRCA1 B, and BRCAC, with different associated proteins) that participate in distinct processes to repair DNA damage (Deng and Brodie, 2000; Huen and Chen., 2010, Roy et al., 2011, Scully et al., 1997; Scully et al., 1999; Scully and Livingston, 2000).

Specifically, the BRCA1 A-Complex consists of BRCA1 in association with RAP80, the deubiquitinating (DUB) enzymes BRCC36 and BRCC45, MERIT-40, and the adaptor protein Abraxas 1 (Harris and Khanna., 2011; Her et al., 2016; Savage et al., 2015). The BRCA1 A-Complex participates in DNA repair by targeting BRCA1 to ionizing radiation (IR) inducible foci; this occurs when RAP80 interacts with K63 poly-ubiquitin chains at sites of double strand breaks (DSBs) where the DNA damage marker γH2AX is phosphorylated (Yan and Jetten., 2008). RAP80 ubiquitylation is performed by the E3 ubiquitin ligase, RNF 168/Ubc13, and RNF8, which are targeted to DSBs by MDC1 (Her et al., 2016). BRCA1 is also bound to BRCA1 associated Ring Domain 1 (BARD1), an interaction that is necessary for BRCA1 protein stability, nuclear localization, and E3 ubiquitin ligase activity (Irminger-Finger et al., 2016). In addition, BRCA1 is also a nuclear-cytoplasmic shuttling protein, and increasing evidence suggests that BRCA1 function can be controlled via active shuttling between subcellular compartments (Fabbro et al., 2002; Feng et al; 2004).

We identify a novel tumor-suppressive function for cavin3 mediated through its interaction with BRCA1 leading to regulation of BRCA1 levels, subcellular location, and function. We show that cavin3 controls BRCA1 functions in UV-induced apoptosis and cell protection against DNA damage through downregulation and abolishment of the recruitment of the BRCA1 A-complex to DNA lesions in response to UV damage.

## Results

### Global proteome analyses of cavin3 function reveals a prominent role in DNA repair

As a first step to investigate the cell biology of cavin3 we undertook an unbiased approach to characterize its cellular proteome, using label-free (LFQ) quantitative proteomics. We deleted cavin3 by genome editing in HeLa cells, a well-characterized model system that has been used extensively to study caveolae (Bohmer et al., 2015; Boucrot et al., 2011; Hao et al., 2012; Hirama et al., 2017; Pang et al., 2004; Rejman et al., 2005; Sinha et al., 2011) (**Figure 1A, Supplementary Figure 1A**). Global proteome analyses were carried out with three replicates from matched WT and cavin3 KO HeLa cells. Cells were SILAC-labelled and subjected to mass spectrometric analysis after lysis. Relative protein expression differences were then determined using label free quantitation **(Figure 1A).** A total of 4206 proteins were robustly quantified detected with >2 unique peptides and an FDR <1.0 % in at least 2 out of 3 replicates **(Figure 1A, details** in **Supplementary Table 1**). To validate these results, we immunoblotted for several proteins involved in diverse cellular processes. Levels of these proteins were consistent with the proteomic analysis **(Supplementary Figure 1B)** and their levels were restored by expression of exogenous cavin3, confirming the specificity of the KO effect **(Supplementary Figure 1C)**.

**Figure 1.**
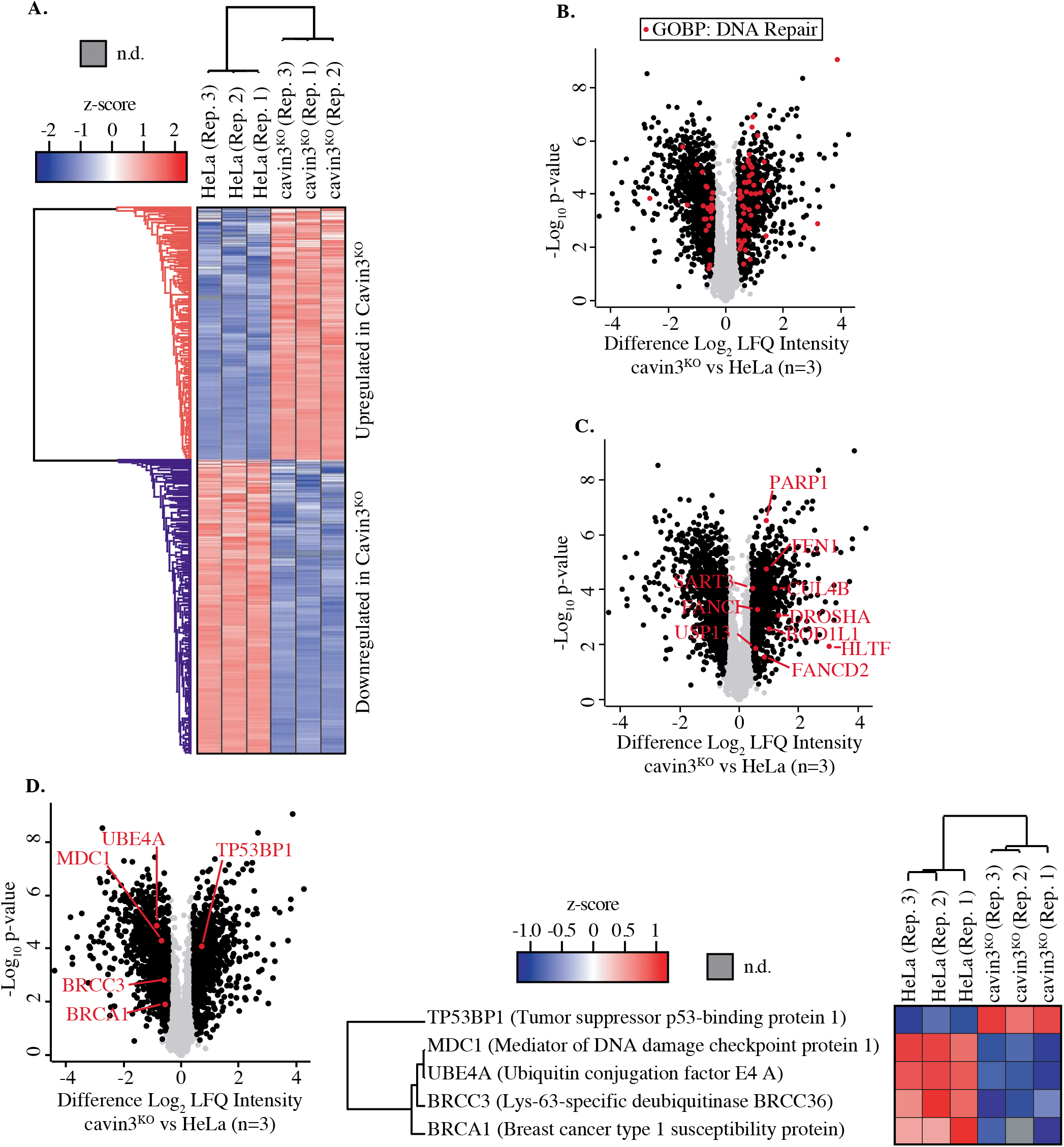
Global proteome analysis of cavin3 KO HeLa cells by label-free quantitative proteomics. (**A**). Z-score for HeLa WT and cavin3 KO cells (Replicates Rep. 1-3) showing upregulated proteins (red) and downregulated proteins (blue). (**B**). Volcano plot showing proteins (red dots) identified by GOBP involved in DNA repair. (**C**). Volcano plot showing DNA repair proteins upregulated in cavin3 KO cells. (**D**). Volcano plot showing proteins of the BRCA1 A-complex, BRCA1, BRCC3, MDC1 and UBE4A downregulated in cavin3 HeLa KO cells and upregulation of TP53BP1 with a heatmap analysis of the expression of each of these proteins in Replicate (Rep. 1-3) HeLa WT and cav in3 KO cells.

Our analysis revealed distinct cavin3-dependent protein networks that might yield new insights into its cellular function. Initial inspection of differentially expressed protein by Gene Ontology analysis revealed that many proteins involved in DNA repair were altered in cavin3 KO cells (**Figure 1B and C and Supplementary Table 2,** see Supplementary Discussion for further analysis of cavin3-dependent pathways). Strikingly, BRCA1 (~1.5 fold decrease) and many components of the BRCA1 A-complex, BRCC36 (~1.5 fold decrease), MDC1 ((~1.7 fold decrease) and the newly described UBE4A (~2.2 fold decrease, Baranes-Bachar et al., 2018) were reduced in cavin3 KO cells that were confirmed by western analysis (**Figure 1D and Supplementary Figure 1D).** In contrast, TP53BP1 was increased in cavin3 KO cells **(Figure 1D and Supplementary Figure 1D).** Accordingly, we elected to pursue the relationship between cavin3 and BRCA1 in greater detail.

### Cavin3 interacts with BRCA1 in vitro and in a model cell system

First, we asked whether cavin3 and BRCA1 might interact in the cytosol. Recent studies suggest that the release of cavin proteins into the cytosol can allow interaction with intracellular targets (Gambin et al., 2014; McMahon et al, 2019; Sinha et al., 2011). To test whether non-caveolar cavin interacts with BRCA1, we used MCF7 cells as a model system. These cells lack endogenous CAV1, cavins, and caveolae (Gambin et al, 2014; McMahon et al., 2019) and so expressed cavin proteins are predominantly cytosolic.

BRCA1-GFP was co-expressed in MCF7 cells with exogenous mCherry-tagged cavins-1, 2, 3, and mCherry-CAV1, and interactions between these proteins were measured in cytoplasmic extracts using two-color Single Molecule Coincidence (SMC) detection. The numbers of photons detected in green and red channels were plotted as a function of time where each fluorescent burst was analyzed for the coincidence between the GFP and cherry fluorescence that reflects co-diffusion of at least two proteins with different tags, the total brightness of the burst, indicating the number of proteins present in the oligomer and the burst profile that is determined by the rate of diffusion and reflects the apparent size of the complex (Gambin et al., 2014). This revealed a specific association between BRCA1 and cavin3-mCherry, but not with the other cavin proteins **(Figure 2A-E).** Quantitatively, 60% of BRCA1-GFP associated with cavin3-mCherry **(Figure 2D)**. The distribution of bursts revealed the behavior of monomeric GFP. This data was used to calibrate the brightness profile and estimate the number of BRCA1-GFP molecules. We concluded that over-expressed BRCA1 primarily exists in a dimeric state when expressed in MCF7 cells and that a dimer of over-expressed BRCA1 interacts with a monomer of exogenous cavin3 **(Figure 2F).** Similar results were obtained when BRCA1-GFP and cavin3-mCherry were co-expressed in MDA-MB231 cells, a cell line with endogenous caveolar proteins and abundant caveolae at the plasma membrane **(Supplementary Figure 2A-E).** This implied that BRCA1 and cavin3 can interact in the cytosol, irrespective of the caveolar state of the cells.

**Figure 2.**
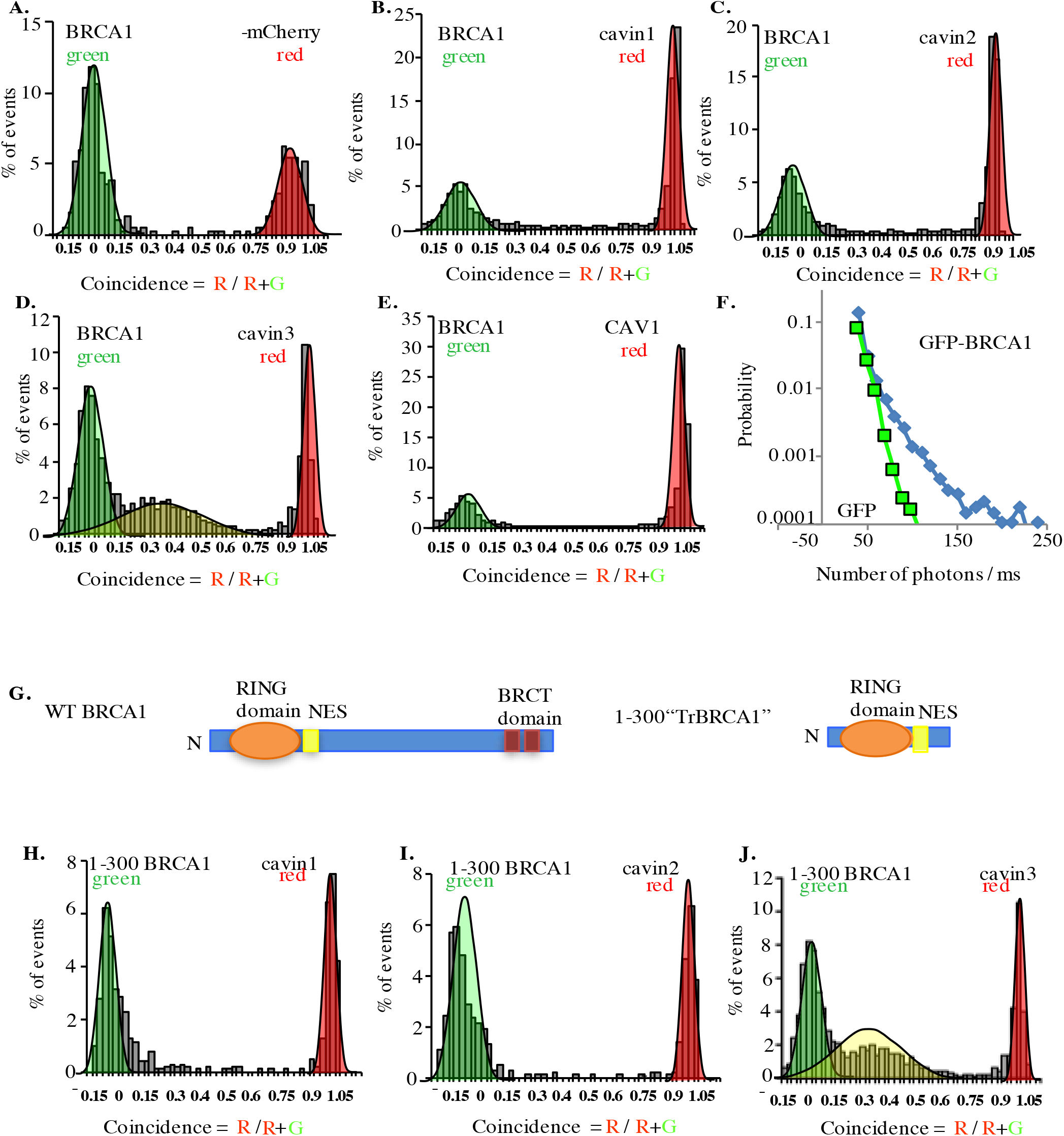
Single molecule analysis of BRCA1 with cavin3-mCherry in MCF-7 cells. **(A).** Two-color single molecule fluorescence coincidence of BRCA1-GFP with **(A).** mCherry control, **(B).** mCherry-cavin1, **(C).** mCherry-cavin2, **(D).** mCherry-cavin3, **(E).** mCherry-CAV1 coexpressed in MCF-7 cells. The green curve represents BRCA1-GFP only events, the red curve represents mCherry only events and the yellow curve represents BRCA1-GFP + Cherry events. **(F).** Distribution of burst brightness measured for BRCA1-GFP (blue) and GFP control (green). **(G).** Schematic representation of domain organization of full-length wildtype (WT) BRCA1 and the truncated (Tr) 1-300 BRCA1 constructs. Nuclear export signal-NES, BRCA1 C Terminus domain (BRCT) domain, N-N Terminus **(H-J).** Two-color single molecule fluorescence coincidence of 1-300 BRCA1 with **(H).** cavin1, **(I).** cavin2 and **(J).** cavin3 expressed in Leishmania cell-free lysates. More than 1000 events were collected in all cases.

We then used a *Leishmania* cell-free system (Gambin et al., 2014; Sierecki et al., 2013) to test whether these proteins can interact directly. Indeed, a construct bearing the first 300 amino acids of BRCA1 (1-300, tr-BRCA1), which contains the nuclear export signal (NES) and BARD1 binding sites **(Figure 2G),** was associated with cavin3 **(Figure 2J)**, but not with the other cavin proteins **(Figure 2H-I)**. These data suggest that cavin3 directly binds to the N-terminus of BRCA1.

Finally, we used in *situ* proximity ligation assay (PLA) technology (Soderberg et al., 2007) to probe for protein-protein association within intact cells. GFP-tagged cavins or CAV1-GFP were expressed in MCF7 cells and potential associations between transgenes and endogenous BRCA1 were analysed using anti-BRCA1 and anti-GFP antibodies. Positive interactions in PLA analyses are revealed by fluorescent puncta **(Figure 3A-E)**. Puncta were clearly evident throughout the cytosol of cells expressing cavin3-GFP, but not with the other cavins, CAV1-GFP or GFP alone (**Figure 3A-E,** quantitation in **Figure 3F**). Additional experiments using different combinations of antibodies (eg. rabbit antibodies against endogenous BRCA1 together with mouse anti-GFP antibodies **(Supplementary Figure 3)** yielded similar results. Control experiments (GFP alone, BRCA1 alone, absence of PLA probes and with no antibody) yielded few puncta **(Supplementary Figure 4A-E).** Taken together, these studies suggest that BRCA1 can interact with cavin3 directly *in vitro*, and that expressed cavin3 can associate with endogenous BRCA1 in cells.

**Figure 3.**
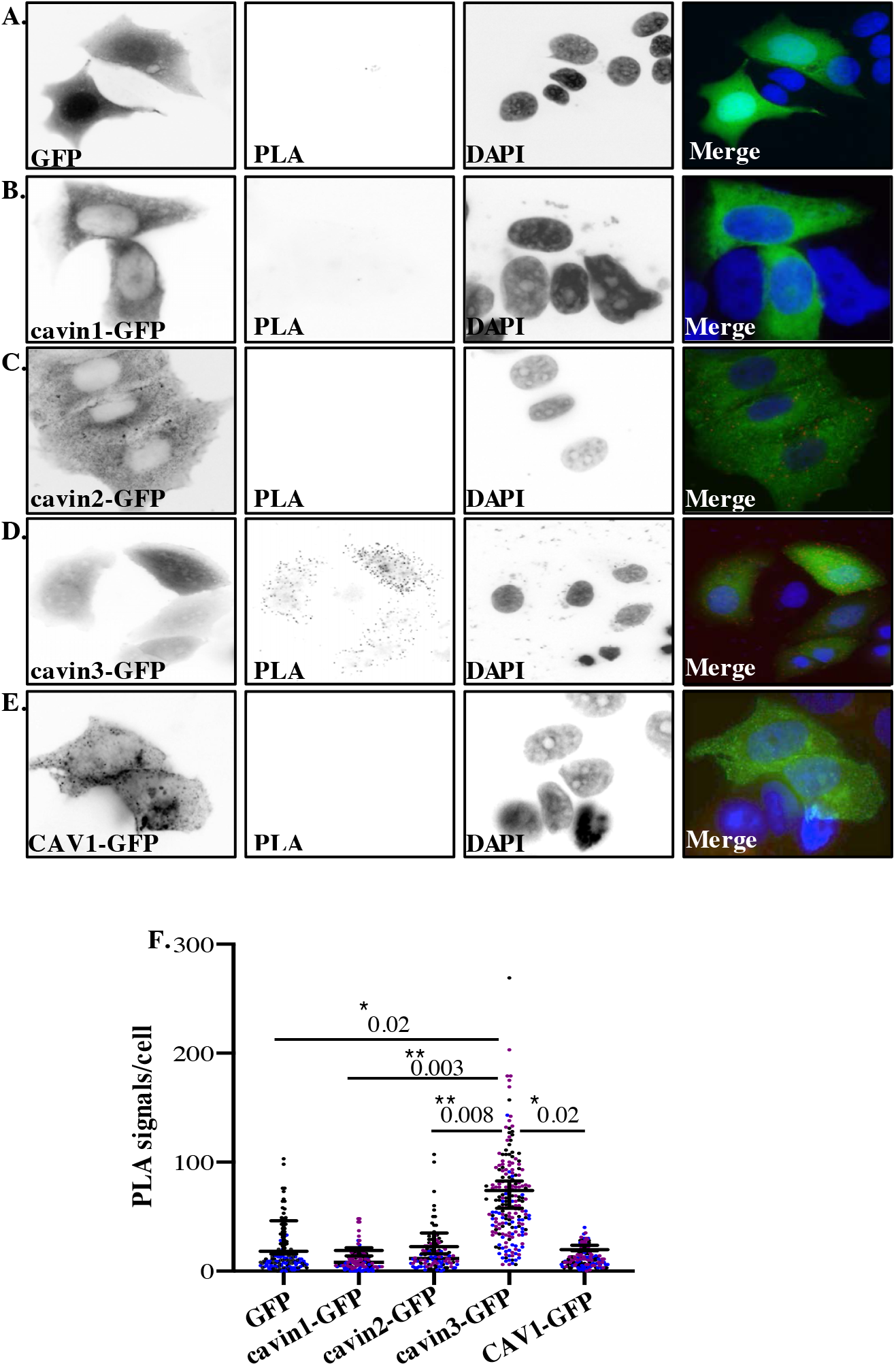
PLA analysis of cavin3 and BRCA1 interaction in MCF7 cells. **(A-E).** Immunofluorescence microscopy in combination with PLA for protein-protein interactions (red dots) within single cells of stably expressing **(A).** MCF7/GFP, **(B).** MCF7/cavin1-GFP, **(C).** MCF7/cavin2-GFP **(D).** MCF7/cavin3-GFP and **(E**). MCF7/CAV1-GFP using monoclonal GFP and polyclonal BRCA1 antibodies. DNA was counterstained with DAPI (blue). Scale bars represent 10 μm. **(F).** Number of red dots/PLA signals in 40-50 cells for each MCF7/GFP expressing cell line was quantified from 3 independent experiments using a nested ANOVA Each biological replicate is color coded and the Mean ± SEM is presented as a black bar. ** p<0.05, ** p<0.01.

### Cavin3 regulates BRCA1 protein expression and localization

We next examined the relationship between cavin3 and the subcellular localization of BRCA**1**. Immunofluorescence revealed a typical nuclear staining pattern for endogenous BRCA1 with little cytoplasmic staining in control MCF7 cells and cells expressing cavin1-GFP **(Figure 4A)**. In contrast, the expression of cavin3-GFP increased cytosolic staining for endogenous BRCA1 **(Figure 4A),** and this was confirmed by quantitative analysis of the protein distribution **(Figure 4B).** Western blotting revealed that cavin3-GFP selective increased total cellular levels of BRCA1 (**Figure 4C**, quantitation in **Supplementary 5A**), and this represents a post-transcriptional effect of cavin3, as BRCA1 mRNA levels were not significantly increased **(Supplementary Figure 5B).** Interestingly, the proteasome inhibitor, MG132, increased BRCA1 levels in control cells, consistent with evidence for proteasomal degradation of BRCA1 (Choudhury et al., 2004) but it did not increase the already-elevated levels of BRCA1 found in cavin3-GFP cells (**Figure 4D**, quantitation in **Supplementary Figure 5C**).

**Figure 4.**
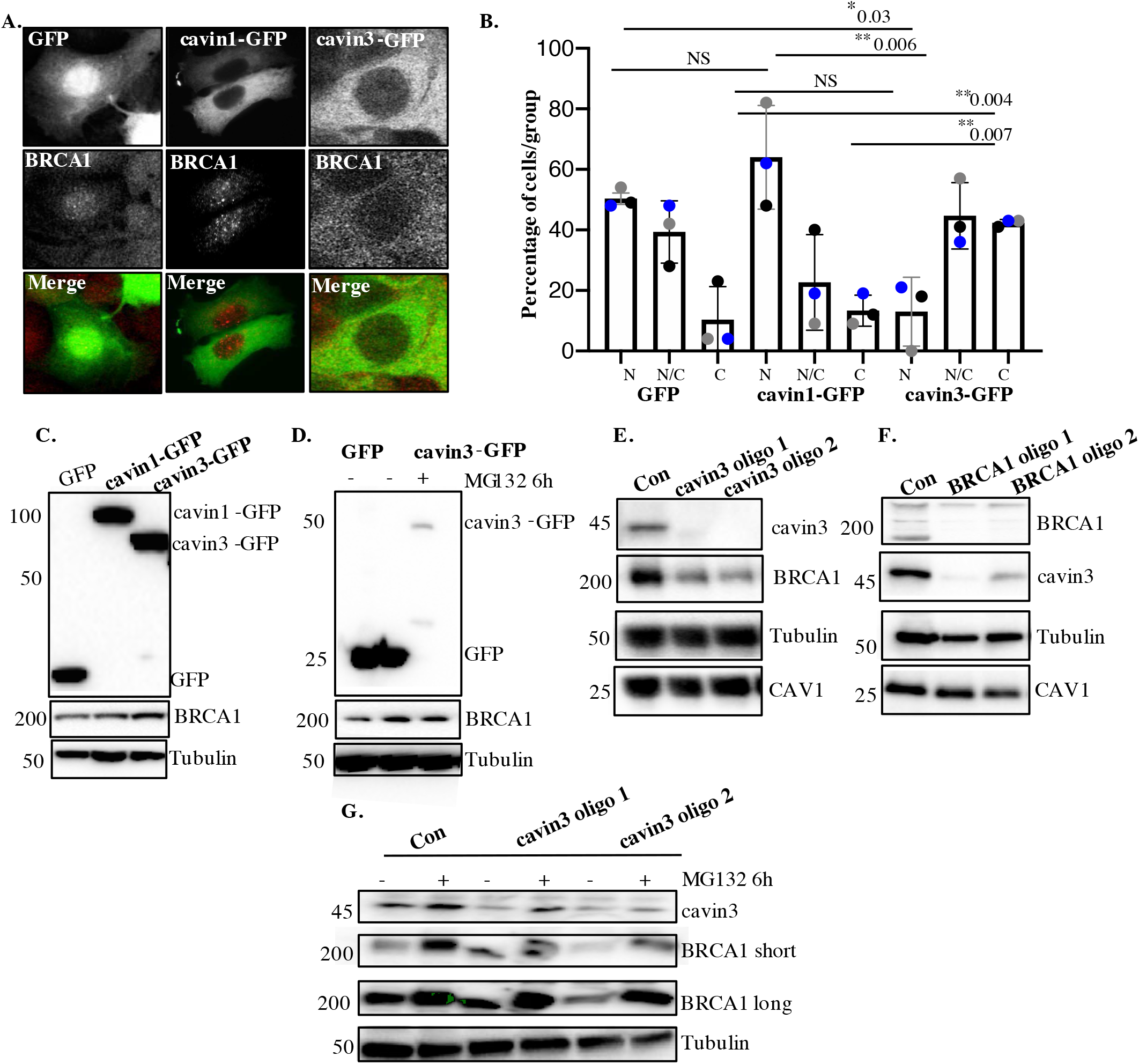
Cavin3 regulates BRCA1 protein expression and localization. **(A).** Representative image of MCF7 cells stably expressing GFP alone, cavin1/GFP and cavin3/GFP fixed and stained with a BRCA1 antibody. **(B).** Percentage of MCF7 cells showing strictly nuclear, nuclear-cytoplasmic or cytoplasmic localization of BRCA1 was counted for 50 cells from 4-5 independent experiments as Mean ± SD using a one-way ANOVA and Bonferroni’s multiple comparisons test. Each biological replicate was color-coded. NS – not significant, * p<0.05, ** p<0.01. **(C)** Lysates from stably expressing MCF7 cells Western blotted for GFP, BRCA1 and Tubulin as a load control. **(D).** MCF7/GFP and MCF7/cavin3-GFP cells, untreated (-) or treated with MG-132 for 6 h. Lysates were Western blotted with GFP, BRCA1, and Tubulin antibodies as a loading control. **(E).** A431 cells treated with control siRNAs (Con) or two siRNAs specific to cavin3. Lysates were Western blotted using cavin3, BRCA1, CAV1 antibodies and Tubulin as the loading control. **(F).** A431 cells treated with control siRNAs or two siRNAs specific to BRCA1. Lysates were Western blotted using cavin3, BRCA1, CAV1 antibodies and Tubulin as the loading control. **(G).** A431 cells treated with Control (Con) or siRNAs specific to cavin3, untreated or treated with MG132 for 6 hours. Lysates were Western blotted using cavin3, BRCA1, and Tubulin as a loading control. Quantitation of all blots in Figure 4 are provided in Supplementary Figure 5A-E.

Dependence of BRCA1 on cavin3 was also evident when cavin3 was depleted in either A431 and MDA-MB231 cells, using two different siRNAs **(Figure 4E,** quantitation in **Supplementary Figure 5D, Supplementary Figure 6A and C).** These cell lines express cavin3, CAV1, and BRCA1 proteins and present caveolae at the plasma membrane **(Supplementary Figure 5E).** In both cases, cavin3 depletion caused a significant decrease in BRCA1 (**Figure 4E,** quantitation in **Supplementary Figure 5D, Supplementary Figure 6A and 6C)** and this was abrogated by proteasome inhibition **(Figure 4G)**. Immunofluorescence staining revealed that BRCA1 was reduced in the cytosol and nuclei of cavin3 siRNA cells **(Supplementary Figure 7)**. Interestingly, depletion of BRCA1 with two independent siRNAs significantly decreased endogenous cavin3 protein levels in these cells (**Figure 4F,** quantitation in **Supplementary Figure 5F, Supplementary Figure 6B and 6D)**. Taken with our earlier work on HeLa cells, these results collectively show that cavin3 can support BRCA1 protein levels in a variety of cancer cell systems.

### Cavin3 associates with BRCA1 when caveolae disassemble

What might induce cavin3 to interact with BRCA1? A variety of stresses cause caveolae to flatten and disassemble, releasing cavins into the cytosol. We, therefore, hypothesized that stimuli that induce caveola disassembly might induce the association of cavin3 with BRCA1.

First, we tested a role for mechanical stress by swelling cells with hypo-osmotic medium. We used A431 cells for these experiments as they have abundant caveolae. The total association between endogenous cavin3 and endogenous BRCA1, and their association in the nucleus, was significantly increased by hypo-osmotic stimulation, as measured by PLA **(Figure 5A)**. No interaction was seen with a range of control proteins including flotillin 1 and the nuclear proteins PCNA and Aurora kinase **(Figure 5B-E)**. This implied that mechanical disassembly of caveolae could promote the association of cavin3 with BRCA1 both in the cytosol and the nucleus.

**Figure 5.**
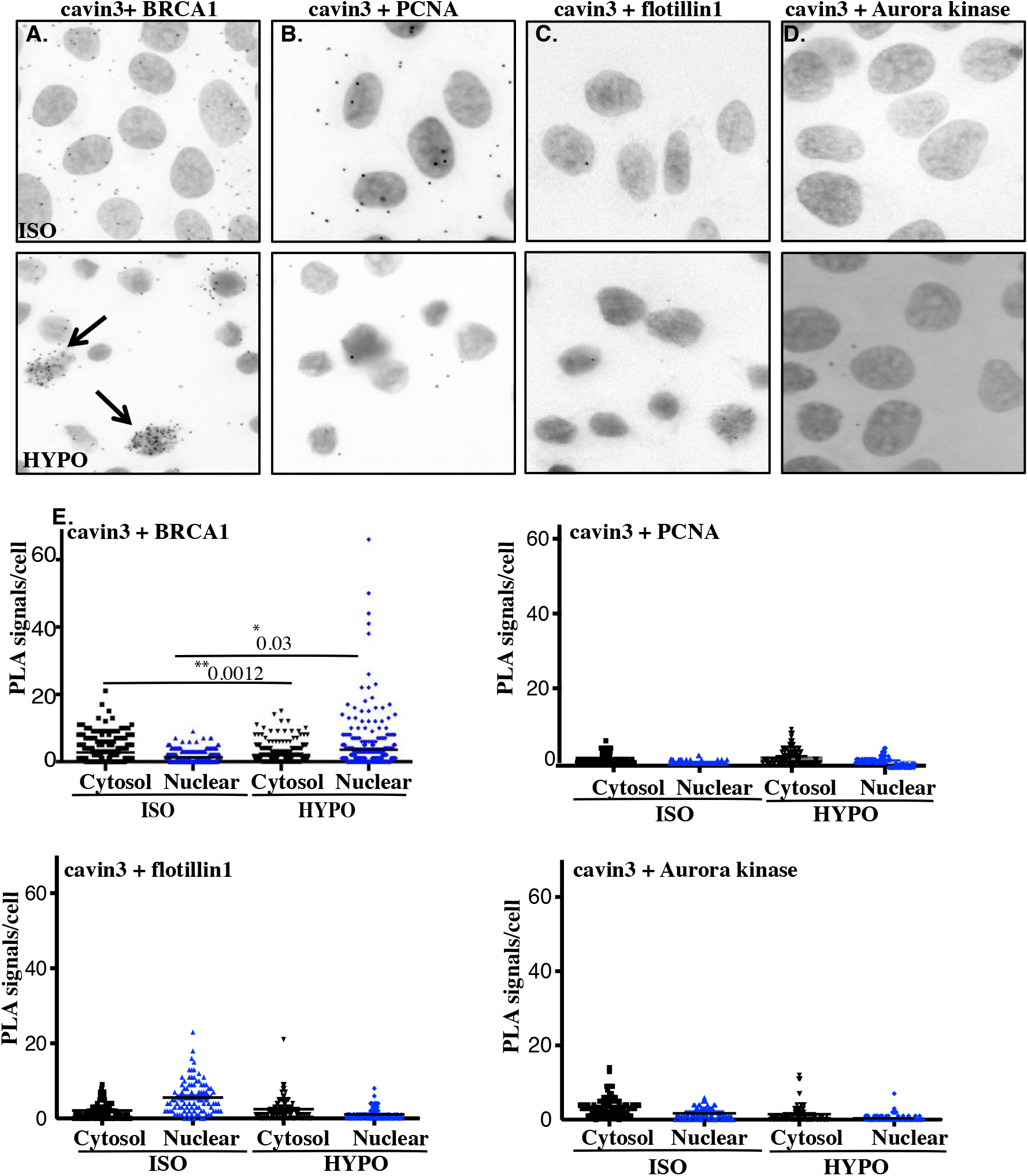
Cellular swelling of A431 cell causes an increase in the BRCA1-cavin3 interaction. **(A).** A431 cells were treated with isotonic (ISO) or hypo-osmotic (HYPO) medium and PLA was performed using cavin3 and BRCA1 **(B).** cavin3 and flotillin1 **(C).** cavin3 and PCNA and **(D).** cavin3 and Aurora kinase antibodies as controls for PLA. DNA was counterstained with DAPI (blue). Scale bars represent 10 μm. **(E).** Total number of PLA signals in the cytosol and the nucleus of cells as defined by DAPI staining in 50 cells for each pair of antibodies quantified from three independent experiments using a nested ANOVA with the Mean ± SEM represented by the black bar, * p<0.05, **p<0.01.

Nest, we tested the effect of non-mechanical stimuli by exposing cells to either UV (2 min pulse, 30 min chase) or oxidative stress with hydrogen peroxide (H_2_O_2_, 200 μM, 30 min). PLA showed that the interaction between endogenous BRCA1 and cavin3 was increased by both these stimuli **(Figure 6A-D top panel,** quantitation in **Figure 6E)**. A more extended time course further demonstrated that association between these proteins was evident at 30 min and maintained at low levels for up to 4 hours **(Figure 6F-G)**. Interestingly, this coincided with a decrease in the interaction between cavin3 and cavin1, which occurs in caveolae **(Figure 6A-D, bottom panel,** quantitation in **Figure 6G).** Similar effects were seen in MDA-MB231 cells **(Supplementary Figure 8).** Control experiments (knockdown of cavin3 or BRCA1 in untreated and UV treated A431 cells) yielded few puncta **(Supplementary Figure 9)**. This was consistent with the notion that cavin3 was moving from caveolae into the cytosol to interact with BRCA1. Our findings indicate that cavin3 can be released to interact with BRCA1 when caveolae disassemble in response to a variety of mechanical and nonmechanical stimuli.

**Figure 6.**
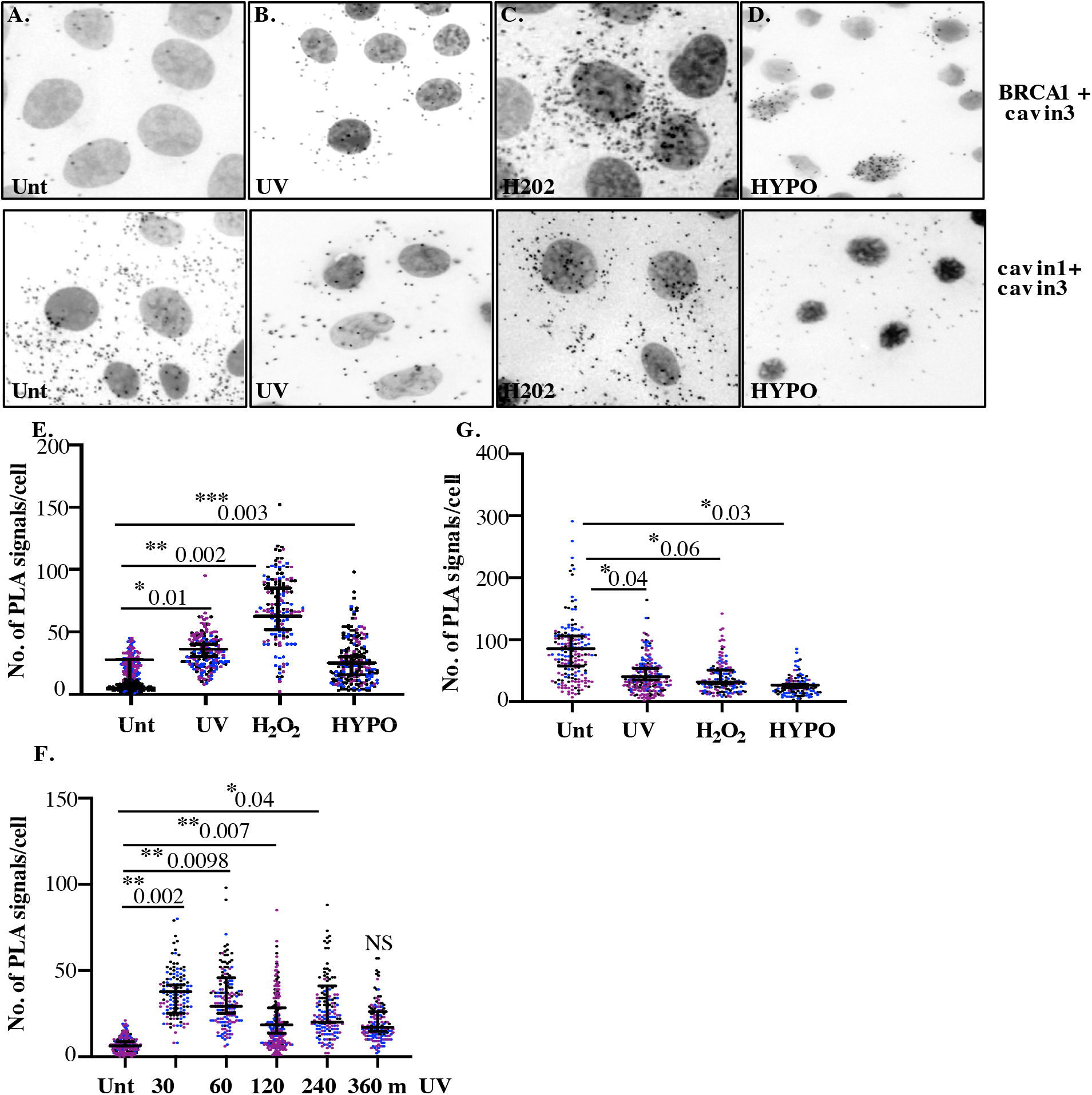
Close association between cavin3 and BRCA1 in A431 cells after stress treatment. **(A).** Immunofluorescence microscopy in combination with PLA visualization of endogenous proteinprotein interactions (red dots) within A431 cells in **(A).** Untreated (Unt.) cells, **(B).** UV treated and a chase time of 30 min, **(C)**. 200 μM H_2_0_2_ (H_2_0_2_) for 30 min and **(D).** Hypo-osmotic treatment (HYPO) for 10 min, top panel–BRCA1 and cavin3 and bottom panel-cavin1 and cavin3. **(E).** PLA signals/cell for cavin3-BRCA1 association in 50 cells/biological replicate with three independent experiments. **(F).** PLA time course analysis after UV treatment and a chase time up to 360 min in 50 cells/biological replicate with three independent experiments. **(G).** PLA signals/cell for cavin1-cavin3 association in 50 cells/biological replicate with three independent experiments. All data was quantified from three independent experiments using a nested ANOVA. Each biological replicate is color coded with the Mean ± SEM presented as a black bar. NS-not significant, * p<0.05, ** p<0.01, ***p<0.001.

### Cavin3 and BRCA1 function similarly in apoptosis in the cytosol and DNA damage sensing in the nucleus

Next, we sought to evaluate the potential functional consequences of this stress-inducible association of cavin3 with BRCA1. As cytoplasmic BRCA1 has been implicated in cell death pathways (Dizin et al., 2008; Thangaraju et al., 2000; Wang et al., 2010), we asked if cavin3 affects the sensitivity of cells to apoptosis induced by UV exposure. We found that LDH release, used as an index of membrane damage, was consistently increased after 2 min UV exposure in MCF7 cells that over-expressed cavin3-GFP, but not with cavin1-GFP **(Figure 7A).** That this cell damage reflected induction of apoptosis was confirmed by staining for annexin V (which marks early apoptosis, **Figure 7B**) and the DNA dye 7-amino-actinomycin 7 (7-AAD, late apoptosis, **Figure 7C**). Both apoptotic markers were enhanced by cavin3-GFP over-expression. Thus, cavin3 could sensitize MCF7 cells to UV-induced apoptosis.

**Figure 7.**
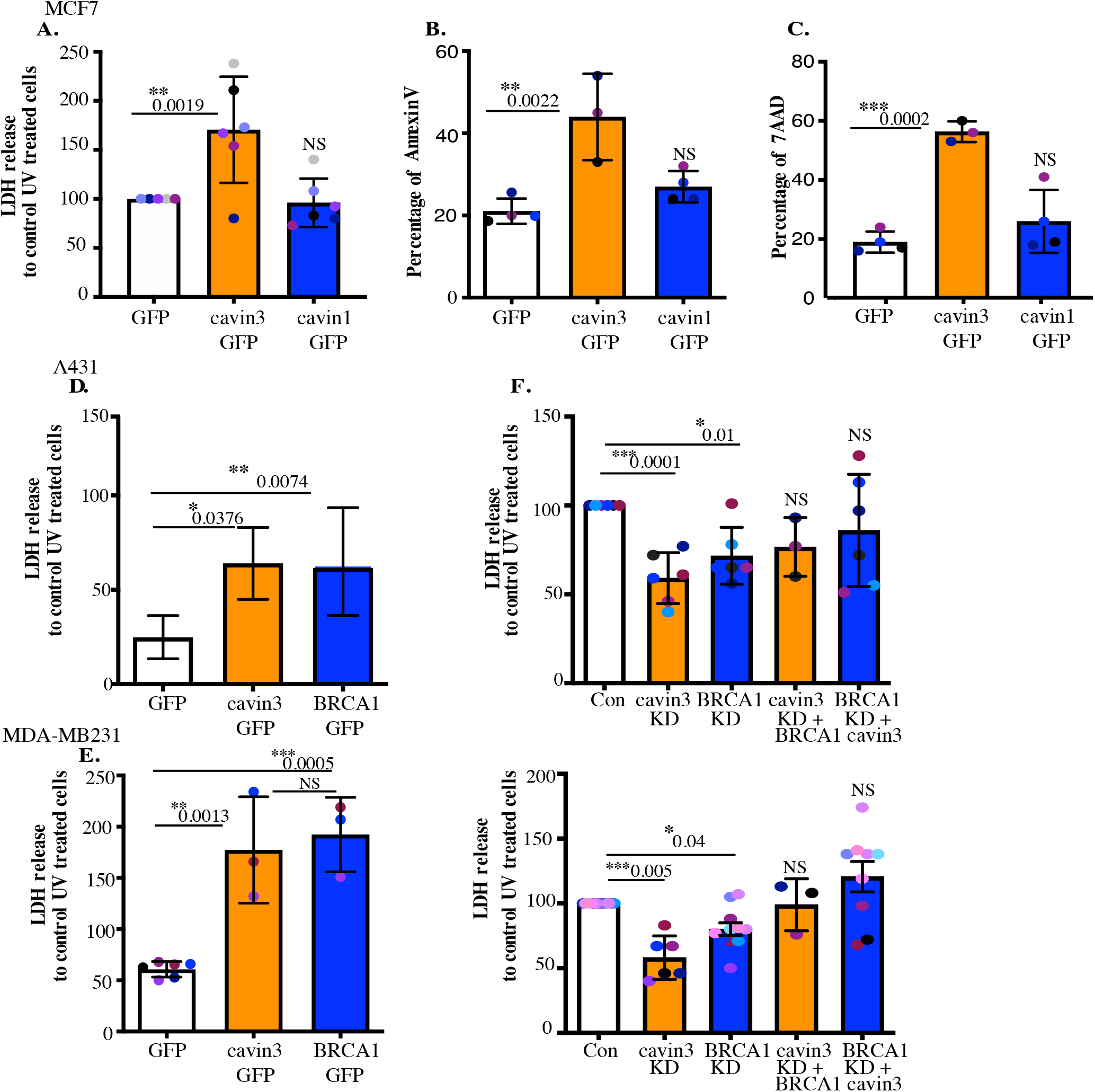
Cavin3 potentiates BRCA1 functions in apoptosis. **(A)** LDH release of MCF7/GFP, cavin3-GFP and cavin1-GFP cells subjected to UV treatment and a 6 h chase. LDH release is expressed as a percentage to control GFP cells from six independent experiments presented as Mean ± SD using a one-way ANOVA and Bonferroni’s multiple comparisons test. **(B).** Annexin V positive cells after UV treatment and a 6 h recovery time in MCF7 cells presented as Mean ± SD using a one-way ANOVA and Bonferroni’s multiple comparisons test from three independent experiments. **(C)** 7-AAD positive cells after UV treatment and a 24 h recovery time in MCF7 cells presented as Mean ± SD using a one-way ANOVA and Bonferroni’s multiple comparisons test from three independent experiments. **(D).** A431 cells and **(E).** MDA-MB231 cells were transfected with GFP, cavin3-GFP or BRCA1-GFP. Results are the relative percentage of LDH release to GFP as Mean ± SD using a one-way ANOVA and Bonferroni’s multiple comparisons test from at least 3 independent experiments. **(F)** A431 cells and **(G).** MDA-MB231 cells were treated with control, cavin3 or BRCA1 specific siRNAs. Cavin3-depleted A431 and MDA-MB231 cells were transfected with BRCA1-GFP for 24 hours. BRCA1 depleted A431 and MDA-MB231 cells were transfected with cavin3-GFP for 24 hours. All cells were UV treated and LDH release was measured and calculated relative to control siRNA UV treated cells. The results represent independent experiments as Mean ± SD using a one-way ANOVA and Bonferroni’s multiple comparisons test from three independent experiments. Each biological replicate is color coded. NS – not significant, * p<0.05, ** p<0.01, ***p<0.001.

We then asked whether this effect also operated in cancer cells with endogenous expression of cavin3. Indeed, over-expression of cavin3-GFP significantly increased LDH release from UV-treated A431 and MDA-MB231 cells **(Figure 7D and 7F)**. Furthermore, depletion of endogenous cavin3 reduced LDH release from these cells after UV stimulation **(Figure 7E and G,** controls in **Supplementary Figure 10A and 10B).** Together, these findings indicate that cavin3 sensitizes cells to apoptosis induced by UV.

BRCA1 also sensitized A431 and MDA-MB231 cells to apoptosis, as evident when exogenous BRCA1 was overexpressed or the endogenous protein was depleted **(Figure 7E and 7G,** controls in **Supplementary Figure 10C and 10D)**. Therefore, we further examined the relationship between BRCA1 and cavin3. Overexpression of BRCA1 in cavin3-depleted A431 or MDA-MB231 cells or overexpression of cavin3 in BRCA1-depleted cells restored UV induced apoptosis to control levels. This indicated that these two proteins have a similar sensitizing effect on UV induced apoptosis (**Figure 7E and 7G)**. These results suggest a pro-apoptotic role for both cavin3 and BRCA1 in stress-induced cancer cells. Similarly, in MCF7 cells expression of cavin3 alone or in combination with BRCA1 restored the sensitivity of BRCA1 KD cells to UV-induced apoptosis **(Supplementary Figure 10E)**. We further exposed WT and cavin3 KO HeLa cells to a range of stressors that allow interaction with BRCA1, including hypo-osmotic medium, UV and oxidative stress (**Supplementary Figure 11A-D**). Cavin3 KO cells exhibited enhanced resistance to all stressors and apart from oxidative stress, this was time-dependent **(Supplementary Figure 11A-D).** Overall, these findings suggest that BRCA1 and cavin3 participate together in the cellular stress response.

### Cavin3 protects against stress-induced DNA damage

In addition to promoting apoptosis, BRCA1 - notably via its BRCA1 A-complex - has also been implicated in DNA repair to limit the mutational risk in stressed cells that evade apoptosis. As noted earlier, we found that BRCA1 A-complex components were reduced at steady-state in cavin3 KO Hela cells **(Figure 1A).** Next, we examined the effect of UV treatment on the level of these components in WT and cavin3 KO HeLa cells. As shown in **Figure 8 A-C,** UV treatment of WT cells upregulated the expression of cavin3, BRCA1, the DNA damage marker, RAD51, and the A-complex proteins MDC1, Rap80, RNF168 and Merit40. Strikingly, the upregulation of BRCA1, RAD51 and the A-complex proteins was dramatically reduced in cavin3 KO cells (**Figure 8A and C,** quantitation in **Supplementary Figure 12**). This suggested that cavin3 can influence the ability of BRCA1 to repair damaged DNA.

**Figure 8.**
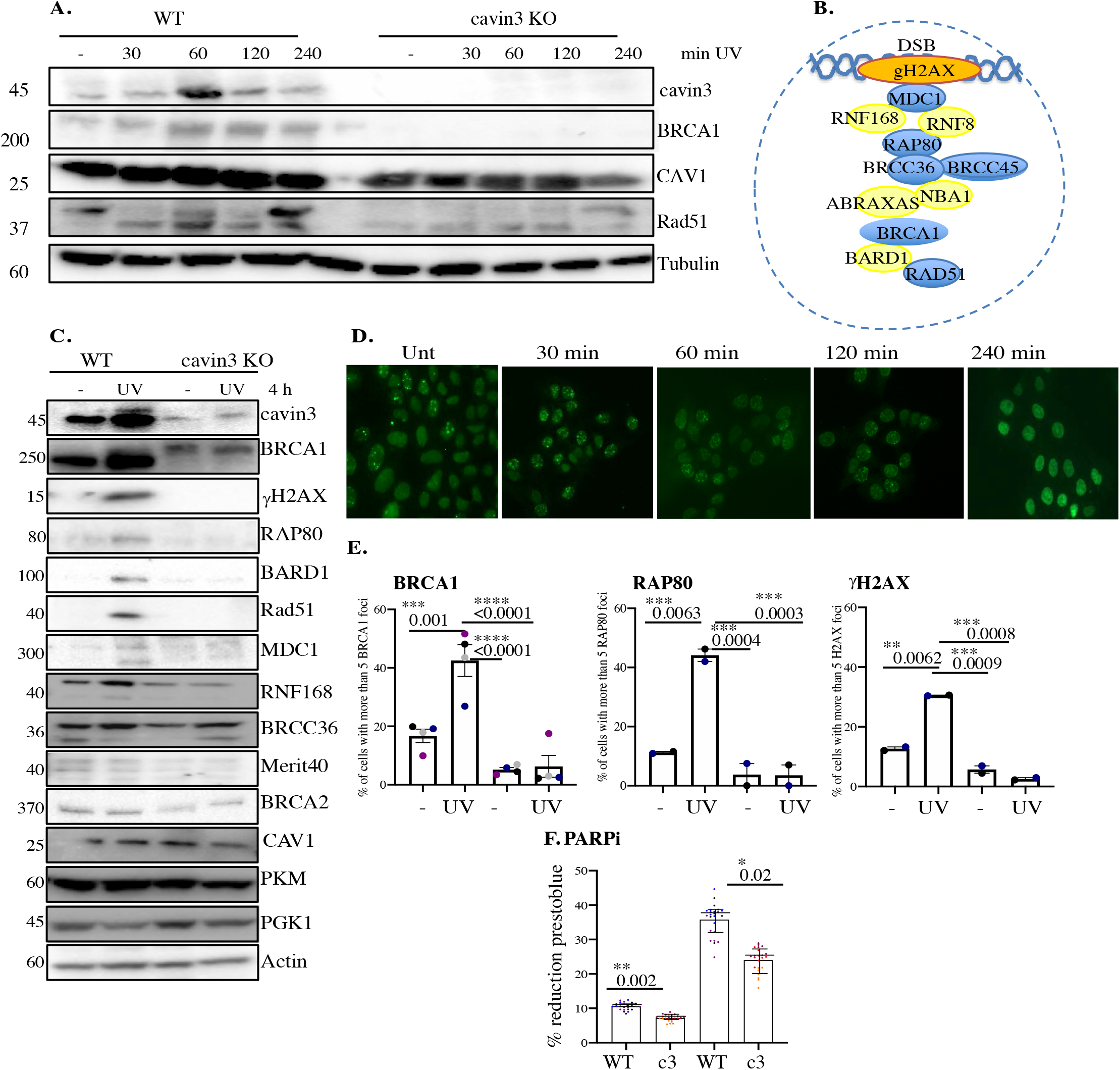
Cavin3 deficient HeLa cells exhibit abolishment of DNA repair. **(A**). Representative Western blot analysis of WT and cavin3 KO cells UV time course for cavin3, BRCA1, CAV1, Rad51, and Tubulin. **(B).** Protein components of the BRCA1 A-complex. Blue colored circles; proteins downregulated in the LFQ proteomics and yellow colored circles; proteins not detected in the LFQ proteomics of cavin3 KO cells. **(C).** Representative Western blot analysis of cavin3, BRCA1, γH2AX, UIM1C/Rap80, BARD1, Rad51, MDC1, RNF168, BRCC36, Merit40, BRCA2, CAV, PKM, PGK1 and Actin in WT and cavin3 KO HeLa cells untreated (-) or UV treated (UV) followed by a 4 hour chase. Quantitation of protein levels from three independent experiments is presented in Supplementary Figure 12. **(D).** Representative immunofluorescence images of BRCA1 foci after UV treatment in WT HeLa cells. **(E).** Percentage of cells with more than five BRCA1 foci, Rap80 foci and γH2AX foci in WT and cavin3 KO cells following UV treatment and a 30 min chase. The results are presented as Mean ± SD using a oneway ANOVA and Bonferroni’s multiple comparisons test from three independent experiments. **(F).** WT and cavin3 KO cells were treated with a PARP inhibitor (PARPi) 50 μ M AZD 2461. Presto blue reagent was added to plates immediately and were read at 570 and 600 nm at 6 hours and 24 hours. The % reduction prestoblue calculated from eight wells/replicate experiment is presented as the Mean ± SEM using a nested ANOVA from three independent experiments. Each biological replicate is color coded. * p<0.05, ** p<0.01.

To test this, we first examined the response of BRCA1 to DNA damage. BRCA1 relocates to form foci at sites of DNA double-strand breaks (DSBs). Indeed, we found that BRCA1 foci increased within 30 min of UV irradiation **(Figure 8D and E),** however this was significantly reduced in cavin3 KO cells **(Figure 8E).** Similarly, the recruitment of RAP80 and γH2AX was compromised in cavin3 KO cells, suggesting that DNA repair might be fundamentally compromised in these cells **(Figure 8E).** To test the functional implications of this result, we evaluated the response of cavin3 KO cells to inhibition of PARP, an ADP-ribosylase that acts as an early responder in DNA repair. PARP inhibition leads to cell death because DSBs cannot be efficiently repaired (reviewed by Noordermeer and Attikum, 2019). Indeed, cell viability studies revealed that cavin3 KO cells were more sensitive to the PARP1 inhibitor AZD2461 than controls **(Figure 8F)**. Together, these results show that cavin3 also promotes BRCA1-mediated DNA repair.

## Discussion

Here we describe a novel role for caveolae and the cavin3 protein in regulating the critical tumor suppressor, BRCA1. Our studies raise the intriguing possibility that by releasing cavins, that can be triggered by mechanical and non-mechanical stimuli such as UV and oxidative stress (McMahon et al., 2019, and this study), that caveolae can act as general sensors and transducers of cellular stress. Our findings suggest that defining the role of the cavin proteins may provide new insights into the functions of caveolae in pathological conditions such as cancer. Cavin3 may represent a promising therapeutic target in breast cancer through its ability to act both inside and outside of caveolae, by modulating specific signaling pathways (Hernandez et al., 2013) and by interacting with and modulating the expression of many proteins such as BRCA1, as shown here, and PP1alpha as previously described (McMahon et al., 2019).

The possibility of an interaction between BRCA1 and cavin3 was first suggested some 15 years ago, yet, no experimental evidence to support this interaction has been published to date. Our results provide the first clear evidence that cavin3 directly interacts with BRCA1 and that this occurs when cavin3 is released from caveolae in response to cellular stressors. We established this using multiple techniques, including PLA in MCF7, MDA-MB231 and A431 cells, single-molecule coincidence detection in multiple cancer cell lines (MCF7 and MDA-MB231 cells) and *in vitro* synthesized BRCA1 and cavin3. We were not able to reproducibly coimmunoprecipitate BRCA1 and cavin3. However, this technique can fail to detect weak or transient interactions (Berggard et al., 2007). Instead, the combination of cell-based methods (PLA and single-molecule approaches) and a cell-free direct interaction approach, as used here, provides unequivocal evidence for the proposed interaction between the N-terminus of BRCA1 and cavin3.

We propose that cavin3 can modulate BRCA1 function via multiple mechanisms: direct interaction with the RING domain of BRCA1 **(Figure 2J),** increased localization of BRCA1 to the cytosol **(Figure 4A-B)**, regulation of BRCA1 protein levels **(Figure 4C and 4F, Supplementary Figure 6B)**; modulation of proteasome-mediated protein degradation **(Figure 4G)** and by facilitating the localization of components of the BRCA1-A-complex to sites of DNA lesions in response to UV-induced DNA damage **(Figure 8E)**.

We show that the ubiquitin-proteasomal degradation pathway plays a significant role in the coordinated protein stability of BRCA1 and cavin3 **(Figure 4G)**. Previous studies have identified the RING domain region of BRCA1 as the degron sequence necessary for polyubiquitination and proteasome-mediated protein degradation, which coincides with the interaction domain of BRCA1 identified here for cavin3 (Lu et al., 2007). Our data further supports studies that that the ubiquitin-proteasome plays an important role in regulating BRCA1 during genotoxic stress (Liu et al., 2010). Interaction of BRCA1 with BARD1 protein reduces proteasome-sensitive ubiquitination and stabilization of BRCA1 expression (Choudhury et al., 2004). BARD1 levels were downregulated in cavin3 KO cells **(Supplementary Figure 1D).** Downregulation of BARD1 would be expected to impair BRCA1 function further in cavin3 KO cells as this interaction stabilizes both proteins with a further significant role in homologous recombination (Xia et al., 2003). Further experiments are required to determine if cavin3 disrupts the interaction between BRCA1 and BARD1 and the contribution of BARD1 to the loss of BRCA1 stability and function in these cells.

In addition to its expression, BRCA1 subcellular localization is a significant contributor to its cellular functions (Henderson et al., 2012). Our findings imply that cavin3 may play a role in the cytosolic translocation of BRCA1 (**Figure 4A-B).** It is intriguing to hypothesize that BRCA1 together with cavin3, executes its tumor suppressor function by its critical role in DNA repair in the nucleus and through signaling pathways and interactions that induce the apoptotic machinery in the cytoplasm. This implies that failed repair of DNA damage in the nucleus is linked to the induction of cell death processes and the elimination of damaged cells in the cytosol and that the BRCA1-cavin3 may contribute to this pathway. Interestingly, cells expressing tr-BRCA1 which was identified here as the BRCA1 domain interacting with cavin3 **(Figure 2J),** has been shown to cause BRCA1 translocation to the cytosol and to possess an enhanced sensitivity to UV (Wang et al., 2010). Ongoing investigations to test this idea may provide further insight into the role of BRCA1 nuclear-cytoplasmic shuttling and determination of cell fate (survival vs. death). Furthermore, these data also point to the potential use of BRCA1 shuttling as a novel therapeutic strategy by which manipulation of BRCA1 localization can control cellular function and sensitivity to therapy.

Cavin3 KO HeLa cells exhibited abolished recruitment of the BRCA1 A-complex to UV-induced DNA damage foci such that BRCA1-dependent DNA damage repair was compromised in these cells **(Figure 8E)**. This was further correlated with a decrease in the protein levels of the components of the BRCA1 A-complex, specifically in these cells **(Figure 8D).** This is consistent with the observation that the loss of any member of the RAP80-BRCA1 complex eliminates observable BRCA1 foci formation, as the BRCA1 A-complex requires all its protein components to be stable to optimally recruit BRCA1 to DSBs (Jiang and Greenberg, 2015). Recent studies from our laboratory have shown that γH2AX phosphorylation is compromised in cavin3 KD cells and that γH2AX forms a complex with the protein phosphatase PP1alpha, whose activity was regulated by cavin3 (McMahon et al, 2019). γH2AX is one of the initial factors that recruit checkpoint and DNA repair proteins to DSBs. Failure of cavin3 KO cells to phosphorylate H2AX may further compromise DNA repair mechanisms in these cells.

In addition, LFQ whole proteomics revealed that cavin3-deficient cells phenotypically resemble BRCA1-deficient cells in exhibiting impaired homologous recombination with significant downregulation of components of the BRCA1 A-complex and sensitivity to PARP inhibition (**Figure 8**). The LFQ proteomics revealed that cavin3 KO cells upregulate many proteins involved in the protection and maintenance of the replication fork and postreplication repair, suggesting that deficiency of cavin3 regulates pathways similar to BRCA1 (**Figure 1, Supplementary Table 1).** These pathways collectively may account for many of the characteristic features of genomic instability in the familial breast, and ovarian cancers and cavin3 KO cells provide an alternative model cell line for further investigation. These findings support further evaluation of replication stress and alternative repair pathways in cavin3 KO cells (see Supplementary Discussion for further analysis of cavin3-dependent pathways).

Finally, the example of cavin3 leads us to propose a general model for cell stress sensing mediated by cavins when they are released from caveolae to interact with intracellular targets. Rigorous control of such a pathway would require that cytosolic levels of cavins must be kept low under steady-state conditions. Recent work shows that this can be achieved by ubiquitination of a conserved phosphoinositide-binding patch on cavins that is only exposed when cavins are released from caveolae (Tillu et al., 2015). In the absence of stabilizing interactions, the released cavin protein will undergo proteasomal degradation, but, as shown here, interaction with BRCA1 stabilizes cavin3, preventing degradation. We propose that the interaction of cavin3 with BRCA1 in response to short term stress can facilitate DNA repair. With a prolonged stress this can trigger apoptosis as a protective mechanism. This forms a novel signaling pathway to protect cells against many cellular stresses and represents a new paradigm in cellular signaling that can explain the evolutionary conservation of caveolae and their involvement in multiple signal transduction pathways.

In view of the loss of cavin3 in numerous cancers (Caren et al., 2011; Kim et al., 2014; Lee et al., 2008; Lee et al., 2011; Martinez et al., 2009; Tong et al., 2010; Xu et al., 2001; Zochbauer-Muller et al., 2005) and the crucial role of BRCA1 as a tumor suppressor (King et al., 2003; Miki et al., 1994; Venkitaraman et al., 2002), these studies describing a new functional partner for BRCA1 suggest that cavin3 should be considered in future diagnostic and therapeutic strategies.

## Materials and Methods

### Reagents

Dulbecco’s modified Eagle’s medium (DMEM, Cat no. 10313-021), Z150 L-glutamine 100x (Cat no. 25030-081), Trypsin-EDTA (0.05%) phenol red (Cat no. 25300062) was from Gibco by Life Technologies, Australia. SERANA Foetal bovine serum (FBS), (Cat no. FBS-AU-015, Batch no. 18030416 was from Fisher Biotechnology, Australia). cOmplete™, mini EDTA-free protease inhibitor cocktail (Cat no. 11836170001), PhosSTOP Phosphatase Inhibitors (Cat no. 4906837001), hydrogen peroxide 30% (w/w) solution (Cat no. H1009), AZD2461 (Cat no. SML 1858) and MG132 (Z-Leu-Leu-Leu-al, Cat no. C2211) were from Sigma Aldrich.

### Antibodies

The following antibodies were used: rabbit anti-CAV1 (Cat no. 610060, BD Biosciences, Franklin Lakes, New Jersey, USA), rabbit anti-cavin1 antibody were raised as described previously and was used for immunofluorescence (Bastiani et al., 2009), rabbit anti-cavin3 (Cat no. 16250-1-AP, 1:2000 WB, 1:300 PLA, Millennium Science, Pty, Ltd), mouse anti-cavin1 (1:100 PLA, Abmart, China), rabbit anti-cavin1 (Cat no. AV36965, 1:2000 WB, Sigma Aldrich), mouse anti-cavin3 (Cat no. ABNOH00112464-MO4, 1:200 PLA, VWR International Pty, Australia), rabbit anti-BRCA1 C-20 (SC-642, 1:500 WB, 1:100 IF/PLA, Bio-Strategy Laboratory Products), mouse anti-BRCA1 MS110 (ab16780, 1:1000 WB, 1:100 IF/PLA, Abcam, Australia), rabbit anti-BRCA1 (07-434, 1:1000 WB, Merck Australia), mouse anti-BRCA1 antibody (D-9, SC-6954, 1:50 IF, Bio-Strategy Laboratory Products), rabbit anti-BRCA1 polyclonal antibody (Cat no. PTG-22363-1-AP, 1:1000 WB, Millennium Science, Pty, Ltd), mouse anti-BARD1 antibody (E-11) (Cat no. sc-74559, 1:500 WB, Bio-Strategy Laboratory Products), rabbit anti-BRCA2 antibody (Cat no. 3675-30T, WB 1: 2000, Sapphire Bioscience), mouse anti-GFP (Cat no. 11814460001, 1:300 PLA, 1:4000 WB, Roche Diagnostics), mouse anti-Tubulin (Cat no. AB7291, 1:3000, ABCAM Australia, Pty, Ltd), mouse anti-Flotillin-1 (Cat no. 610821, 1:100 PLA, BD Biosciences), mouse anti-PCNA (Cat no. NA03T, 1:100 PLA, Merck Millipore), mouse anti-Aurora kinase 1 (Cat no. 611082, 1:100 PLA, BD Biosciences), mouse anti-Actin (MAB1051, 1:5000 WB, Merck), rabbit anti-HLTF (Cat no. PTG-14286-1-AP, 1:2000 WB, Millennium Science), rabbit anti-RNF-168 (Cat no. GTX118147, GeneTex), rabbit anti-BRCC36 (Cat no 4311, 1:1000 WB, ProScience), rabbit anti-MDC1 (Cat no. NOVNB10056657SS, 1:100 WB, Novus), rabbit anti-RAP80 (Cat no.14466S, 1:1000 WB, 1:100 IF, D1T6Q rabbit mAb), mouse anti-Rad51 antibody (Cat no. 14B4, WB 1:1000, NOVNB10014825UL, Novus), sheep anti-Merit40 antibody (Cat no. RDSAF6604SP, WB 1:100, R and D systems), rabbit anti-PGK1 antibody (Cat no. GTX107614, WB 1:3000, Sapphire Bioscience), rabbit anti-PKM antibody (Cat no. GTX107977, WB 1:3000, Sapphire Bioscience), rabbit anti-Histone H2A.X-Chip Grade (Cat no. ab20669, 1:1000 WB, Abcam Australia), rabbit Phospho-Histone H2A.X (Ser 139) (20E3) (Cat no. 9718, 1:500 IF, Cell Signaling), anti-gamma H2AX phosphoS139 CHIP Grade (ab2893, WB: 1:3000, ABCAM). Secondary antibodies for immunofluorescence were Alexa Fluor™ 488 Goat anti-Rabbit IgG (H+L) (Cat no. A11034, 1:500, ThermoFisher Scientific), Alexa Fluor™ 546 Goat anti-Mouse IgG (H+L) (Cat no. A11030, 1:500, ThermoFisher Scientific), Alexa Fluor™ 594 Donkey anti-Rabbit IgG (H+L) (Cat no. A-21207, 1:500, ThermoFisher Scientific) and Alexa Fluor™ 594 goat anti-mouse IgG (H+L) (Cat no. A-21203, 1:500, ThermoFisher Scientific). Secondary antibodies for Western blotting were Goat anti-Rabbit IgG (H+L) cross adsorbed secondary antibody, HRP(Cat no. G-21234, 1:5000, Life Technologies), Goat anti-Mouse IgG (H+L) cross adsorbed secondary antibody, HRP (Cat no. G-21040, 1:5000, Life Technologies), Rabbit anti-Sheep IgG (H+L) (Cat no. ab6747, 1:2000, ABCAM).

### Cell Culture

MCF7 cells a human adenocarcinoma cell line with a low invasive phenotype (ATTC HBT-22) were subjected to STR profiling (QIMR Berghofer Cancer Research Institute). MDA-MB231 cells (ATCC HTB-26) a human adenocarcinoma cell line and A431 cells (ATCC CRL-1555), HeLa cells and HeLa KO for cavin3 were cultured in DMEM supplemented with 10% (vol/vol) FBS, 100 units/ml penicillin and 100 μg/ml streptomycin. All cell lines were routinely tested for mycoplasma. MCF-7 cells were seeded at 1 x 106 cells and were transfected with 5 μg pEGFP DNA, pEGFP-cavin1, pEGFP-cavin2, pEGFP-cavin3 or pEGFP-CAV1 DNA using Lipofectamine 2000 (Invitrogen) according to the manufacturer’s instructions. G418 (Sigma Aldrich) was used as a selection drug at 500 μg/ ml.

### Generation of CRISPR cavin3 Knock-out cell lines

The HeLa cavin3 KO cell line was generated as follows according to the protocol published previously (Stroud et al., 2016). Targeting was to the first exon at the second in frame ATG about one-third through the exon as this was easy for targeting.

*Zifit input (in-frame ATGs, **target site**):*

### CAVIN3

GGGGCCTGTGCCCGAGGCGCCGGCGGGGGGTCCCGTGCACGCCGTGACGGTGGTGAC CCTGCTGGAGAAGCTGGCCTCC**ATG**CTGGGAGACTCTGCGGGAGCGGCAGGGAGGCC TGGCTCGAAGGCAGGGAGGCCTGGCAGGGTCCGTGCGCCGCATCCAGAGCGGCCTGGGCGCTCTGAGTCGCAGCCACG

### Zifit output

### TALENs

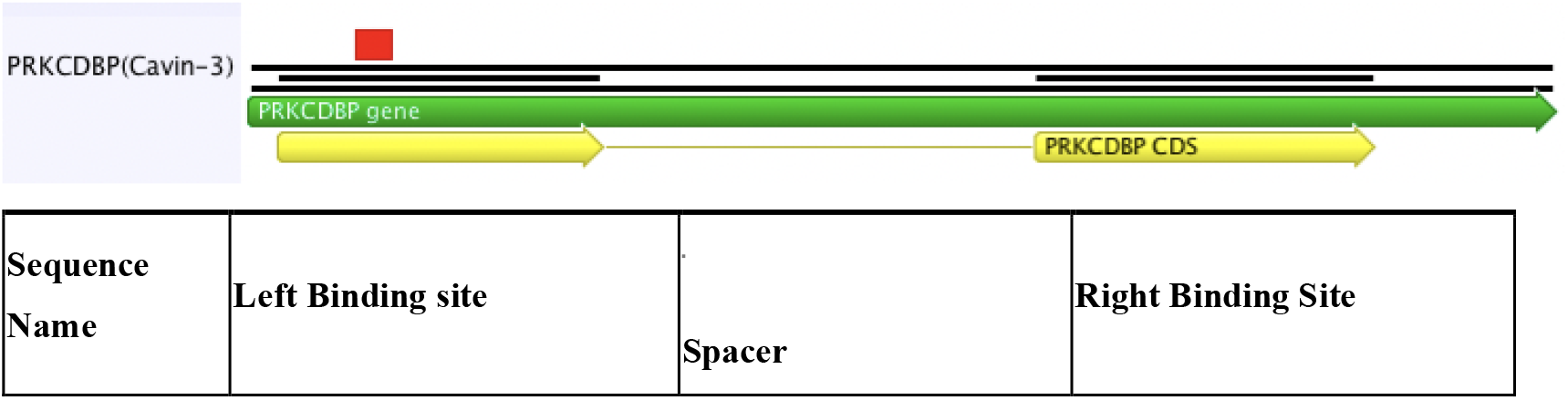
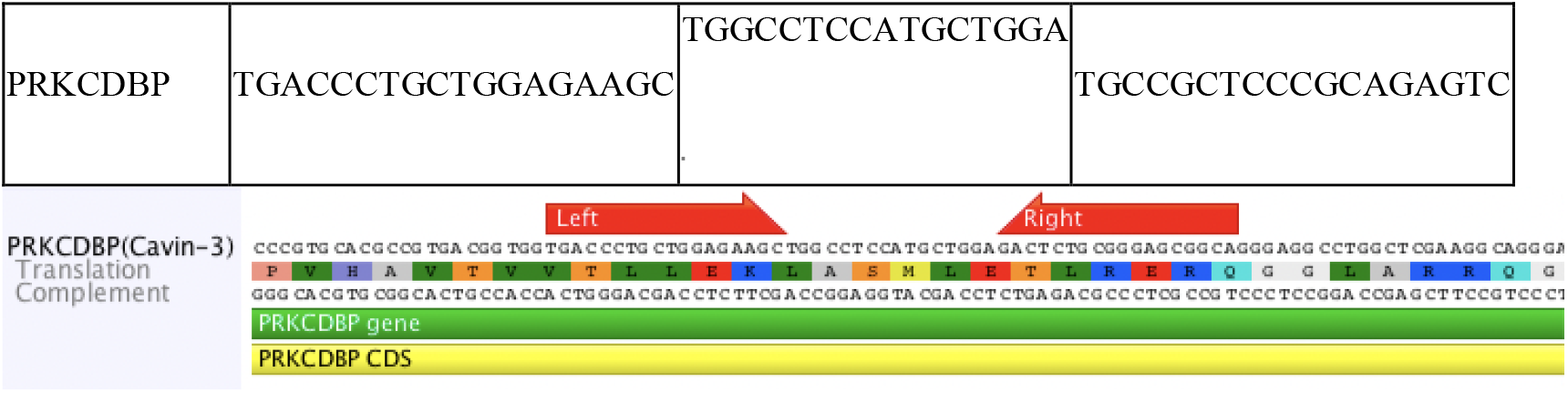

Clonal cells were isolated by dilution into a 96 well plate. Total extract of single clones were prepared and analyzed by Western blotting using rabbit polyclonal anti-cavin3 antibody (Millennium Science). Total deletion of cavin3 was verified by PCR and Western analysis (**Supplementary Figure 1A and 1B**).

### Immunofluorescence

In brief, MCF7, MDA-MB231 and A431 cells seeded onto glass coverslips at 70 % confluence were washed once in PBS and were then fixed in 4 % (vol/vol) PFA in PBS for 20 min at RT. Coverslips were washed three times in excess PBS and were permeabilised in 0.1% (vol/vol) Triton X-100 in PBS for 7 min and blocked in 1% (vol/vol) BSA (Sigma-Aldrich) in PBS for 30 min at RT. The primary antibodies were diluted in 1% (vol/vol) BSA in PBS and incubated for 1 h at RT. Secondary antibodies (Molecular Probes) were diluted in 1% (vol/vol) BSA in PBS and incubated for 1 h at RT. Washes were performed in PBS. Coverslips were rinsed in distilled water and mounted in Mowiol (Mowiol 488, Hoechst AG) in 0.2 M Tris-HCl pH 8.5). The images were taken on a laserscanning microscope (LSM 510 META, Carl Zeiss, Inc) using a 63 X oil lens, NA 1.4. Adjustments of brightness and contrast were applied using Image J software (NIH). The LUT of images for PLA were inverted for better visualization of PLA dots in cells.

### Foci Immunofluorescence

HeLa WT and cavin3 KO cells were pre permeabilized with CSK buffer (10 mM HEPES, 100 mM NaCl, 300 mM sucrose, 3 mM MgCl_2_, 0.7% Triton X-100) for 5 min and were then fixed with 4% PFA/PBS for 15 min, permeabilized with 0.5% Triton X-100 solution for 15 min followed by blocking for 1 h at RT. Cells were then immunostained with primary antibodies against mouse BRCA1 alone (1:40, D9, Santa Cruz), Rap80 alone (1:50, D1T6Q, Cell Signaling), γH2AX alone (1:100, 20669, Abcam) and the appropriate Alexa Fluor™ 488 Goat anti-Rabbit IgG (H+L) (Cat no. A11034, 1:500, ThermoFisher Scientific) conjugated secondary antibodies. Images were taken with a Zeiss microscope. Quantification of the percent of cells was based on foci formation (more than 5 foci/nucleus) was determined from more than 500 cells/experimental condition from two-three independent experiments using an automated plugin for Image J.

### Proximity Ligation Assay (PLA)

Detection of an interaction between BRCA1 and the cavin or CAV1 proteins was assessed using the Duolink™ II Detection Kit (Sigma Aldrich) according to the manufacturer’s specifications. The Duolink™ In situ PLA^®^ Probe Anti-Rabbit MINUS (DUO92005) and Duolink™ In situ PLA^®^ Probe anti-Mouse PLUS (DUO92001) and Duolink™ In situ detection reagents Orange (DUO 92007) were used in all PLA experiments. The primary antibodies used were mouse monoclonal GFP (1:500, Roche) and rabbit polyclonal BRCA1 (1:200, Santa Cruz), cavin3 (1:150, Millennium Science) and PCNA (1:100, Becton Dickinson), cavin3 (1:150, Millennium Science) and Aurora Kinase 1 (1:100, BD Biosciences), cavin3 (1:150 Millennium Science) and Flotillin (1:100, BD Biosciences) and cavin3 (1:150, Millennium Science) and cavin1 (1:100, Abmart). The signal was visualized as a distinct fluorescent spot and was captured on an Olympus BX-51 upright Fluorescence Microscope. The number of PLA signals in a cell was quantified in Image J using a Maximum Entropy Threshold and Particle Analysis where 50 cells in each treatment group were analyzed from at least three independent experiments.

### SDS PAGE and Western blot analysis

For SDS-PAGE, cells were harvested, rinsed in PBS and were lysed in lysis buffer containing 50 mM Tris pH 7.5, 150 mM NaCl, 5 mM EDTA pH 8.0, 1% Triton X-100 with protease and phosphatase inhibitors. Lysates were collected by scraping and cleared by centrifugation at 4 °C. The protein content of all extracts was determined using the Pierce BCA Protein Assay Kit (Cat no. 23225, ThermoFisher Scientific) using bovine serum albumin (BSA) as the standard. Thirty micrograms of cellular protein were resolved by 10 % SDS PAGE and were transferred to Immobilin P 0.45 μm PVDF membrane (Merck). Bound IgG was visualised with horseradish peroxidase-conjugated secondary antibodies and the Clarity™ Western ECL Substrate (Cat no. 1705061, Bio-Rad, Gladesville, New South Wales, Australia).

### Stress Experiments

A431 or MDA-MB231 cells were plated on coverslips at 70% confluency. Cells were either left untreated or were treated with 200 μM H_2_O_2_ for 30 min, 90% hypo-osmotic media for 10 min, or UV treatment for 2 min without media with a UV germicidal light source (UV-C 254 nm) and allowed to recover for 30 min in complete cell culture medium as previously described in McMahon et al, 2019. All cells were fixed and processed for cavin3 and BRCA1 or cavin3 and cavin1 using the Proximity Ligation assay as described.

### Prestoblue Cell Viability assays

HeLa WT and cavin3 KO cells were counted using a hemocytometer and seeded into 96-well plate at 1000 cells/well (8 wells for each treatment) in 90 μl medium per well. Cells were either left untreated or were treated with 90% hypo-osmotic media (90% water in DMEM), UV treatment for 2 min without media with a UV germicidal light source (UV-C 254 nm), 200 μM H_2_O_2_ or 50 μM AZD 2461. After stress addition, 10 μl of PrestoBlue™ Viability Reagent (10x) (Absorbance wavelength: 600 nm) (Thermo Fisher Scientific) was added to cells. The Prestoblue reagent was incubated constantly in wells over a time course from 2 h - 24 h. Control wells containing only cell culture media (no cells) was included in triplicate on each plate for background fluorescence calculations. Plates were returned to a 37 °C incubator. Both absorbance values at 570 nm and 600 nm were measured for each plate in a TECAN Infinite 200 Pro reader, where 570 nm was used as the experimental wavelength and 600 nm as normalization wavelength. Raw data was processed to evaluate the percent reduction of PrestoBlue™ reagent for each well by using the following equation referring to the manufacturer’s protocol:

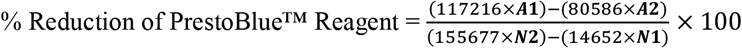

Where: ***A*1**=absorbance of test wells at 570 nm, ***A*2**=absorbance of test wells at 600 nm, ***N*1**=absorbance of media only wells at 570 nm, ***N*2**=absorbance of media only wells at 600 nm.

### RNA Interference

Human cavin3 Stealth siRNAs (set of 3-HSS174185, 150811, 150809) and Human BRCA1 Stealth siRNAs (set of 3 - HSS101089, 186096, 186097) were purchased from Life Technologies Australia Pty Ltd. Two siRNA oligonucleotides to cavin3 or BRCA1 were found to reduce protein levels (oligo 1 and oligo 2) and were transfected into cells at 24 h and 48 h after plating using Lipofectamine 2000 reagent (Invitrogen) with a ratio of 6 μl Lipofectamine to 150 pmol siRNA. Cells were split and harvested after 72-96 h for further analysis.

### Apoptosis Assay

Equal numbers of subconfluent MCF7 cells expressing GFP alone, cavin1-GFP, cavin2-GFP, cavin3-GFP, and CAV1-GFP were seeded on coverslips. Twenty-four hours later, cells were subject to UV-C exposure for 2 min without media. Complete medium lacking phenol red was added to the cells that were left at 37° C to recover. LDH release assay was measured in triplicate samples from 50 μL of conditioned media expressing cells using the Cytotoxicity Detection KitPLUS(LDH) from Sigma Aldrich according to the manufacturer’s instructions. Post-nuclear supernatant from UV exposure cells was also prepared and was subjected to Western blot analysis with antibodies to BRCA1 (1:500, Santa Cruz), GFP (1:3000, Roche) and Tubulin (1:5000, Abcam). For knockdown experiments of cavin3 and BRCA1, after 72 h of knockdown, cells were left untreated or were further transfected with BRCA1-GFP or cavin3-GFP overnight respectively and were then subjected to UV exposure 2 min and a recovery time of 6 h. LDH release was then measured from the cell supernatant in triplicate as indicated in the respective figure legends.

### Single-molecule spectroscopy

Single molecule spectroscopy was performed. Leishmania cell-free lysates were prepared according to (Kovtun et al., 2011, McMahon et al., 2019, Mureev et al., 2009). Where indicated, MDA-MB231 or MCF7 cells were transiently cotransfected with BRCA1-GFP and mCherry alone as the control, cavin1-Cherry, cavin2-Cherry, cavin3-Cherry or CAV1-Cherry constructs. A PNS fraction from the MDA-MB231 and MCF7 cells was prepared in 1 x PBS with protease and phosphatase inhibitors for analysis. Single molecule coincidence measurements were performed using pairs of tagged proteins to ascertain their interaction. One protein of the pair was tagged with GFP, and the other with mCherry, and both were diluted to single molecule concentrations (~1 nM). Two lasers, with wavelengths of 488 nm and 561 nm, (to excite GFP and mCherry, respectively) were focused to a confocal volume using a 40x/1.2 NA water immersion objective. The fluorescence signal from the fluorophores was collected and separated into two channels with a 565 nm dichroic. The resulting GFP and mCherry signals were measured after passing through a 525/20 nm bandpass and 580 nm long pass filter, respectively. The signal from both channels was recorded simultaneously with a time resolution of 1 ms, and the threshold for positive events was set at 50 photons/ms. The coincidence ratio (C) for each event was calculated as C = mCherry/(GFP+mCherry), after subtracting a 6% leakage of the GFP signal into the mCherry channel. Coincident events corresponded to ~0.25<C<0.75. After normalizing for the total number of events (>1,000 in all cases), a histogram of the C values for the protein pair was fitted with 3 Gaussians, corresponding to signals from solely GFP (green), coincidence (yellow), and solely mCherry (red).

### Quantitative mass spectrometry using HeLa WT and Cavin3 KO cells

Samples were prepared for mass spectrometry and analysed as previously described (Stroud et al., 2016). Briefly, cells were lysed in 1% (w/v) sodium deoxycholate, 100 mM Tris-HCl (pH 8.1), Tris(2-carboxyethy)phosphine (TCEP), 20 mM chloroacetamide and incubated at 99 °C for 10 min. Reduced and alkylated proteins were digested into peptides using trypsin by incubation at 37°C overnight, according to manufacturer’s instructions (Promega). Detergent was removed from the peptides using SDB-RPS stage tips as described (Stroud et al., 2016). Peptides were reconstituted in 0.1% % trifluoroacetic acid (TFA), 2% ACN and analysed by online nano-HPLC/electrospray ionization-MS/MS on a Q Exactive Plus connected to an Ultimate 3000 HPLC (Thermo-Fisher Scientific). Peptides were loaded onto a trap column (Acclaim C18 PepMap nano Trap x 2 cm, 100 μm I.D, 5 μm particle size and 300 Å pore size; ThermoFisher Scientific) at 15 μL/min for 3 min before switching the pre-column in line with the analytical column (Acclaim RSLC C18 PepMap Acclaim RSLC nanocolumn 75 μm x 50 cm, PepMap100 C18, 3 μm particle size 100 Å pore size; ThermoFisher Scientific). The separation of peptides was performed at 250 nL/min using a nonlinear ACN gradient of buffer A (0.1% FA, 2% ACN) and buffer B (0.1% FA, 80% ACN), starting at 2.5% buffer B to 35.4% followed by ramp to 99% over 278 minutes. Data were collected in positive mode using Data Dependent Acquisition using m/z 375 - 1575 as MS scan range, HCD for MS/MS of the 12 most intense ions with z ≥ 2. Other instrument parameters were: MS1 scan at 70,000 resolution (at 200 m/z), MS maximum injection time 54 ms, AGC target 3E6, Normalized collision energy was at 27% energy, Isolation window of 1.8 Da, MS/MS resolution 17,500, MS/MS AGC target of 2E5, MS/MS maximum injection time 54 ms, minimum intensity was set at 2E3 and dynamic exclusion was set to 15 sec. Thermo raw files were processed using the MaxQuant platform, Tyanova et al., 2016) version 1.6.5.0 using default settings for a label-free experiment with the following changes. The search database was the Uniprot human database containing reviewed canonical sequences (June 2019) and a database containing common contaminants. “Match between runs” was enabled with default settings. Maxquant output (proteinGroups.txt) was processed using Perseus (Tyanova et al., 2016) version 1.6.7.0. Briefly, identifications marked “Only identified by site”, “Reverse”, and “Potential Contaminant” were removed along with identifications made using <2 unique peptides. Log2 transformed LFQ Intensity values were grouped into control and knockout groups, each consisting of three replicates. Proteins not quantified in at least two replicates from each group were removed from the analysis. Annotations (Gene Ontology (GO), Biological Process (BP)) were loaded through matching with the majority protein ID. A two-sample, two-sided t-test was performed on the values with significance determined using permutation-based FDR statistics (FDR 5%, S0=1). Enrichment analysis of Gene Ontology Biological Process (GOBP) terms was performed on significantly altered proteins using a significance threshold of 4% FDR.

### Statistical Analyses

Statistical analyses were conducted using Microsoft Excel and Prism (GraphPad). Statistical significance was determined either by two-tailed Student’s t-test, one-way ANOVA using the Bonferroni comparisons test with a 95% confidence interval or nest ANOVA, as indicated in the Figure legends. Significance was calculated where * indicates p<0.05, ** indicates p<0.01, *** indicates p<0.001 and **** indicates p<0.0001.

## Abbreviations

ABRAXAS1: Abraxas1, BRCA1 A complex subunit
ACCA: Acetyl-CoA Carboxylase Alpha
ACLY: ATP Citrate Lyase
Alt-EJ: alternative end-joining
ATM: ATM Serine/Threonine Kinase
ATR: ATR Serine/Threonine Kinase
BRCC36: BRCA1/BRCA2-containing complex subunit 36
BRCC45: BRCA1/BRCA2-containing complex subunit 45
DROSHA: Drospha Ribonuclase III
DSBs: double strand breaks
EGFR: EGF Receptor
FANCD2: Fanconi Anemia Complementation Group D2
HLTF: Helicase Like Transcription Factor
MDC1: Mediator of DNA Damage Checkpoint 1
MERIT-40: Mediator of RAP80 Interactions and Targeting subunit of 40 kDa
PARP1: Poly(ADP-Ribose) Polymerase 1
PCNA: Proliferating Cell Nuclear Antigen
RAP80: Receptor-associated protein 80
RNF8: Ring Finger Protein 8
RNF168: Ring Finger Protein 168
RPA2: Replication Protein A2
SMARCAL1: SWI/SNF Related, Matrix Associated, Actin Dependent Regulator of Chromatin, Subfamily A Like 1
TOPBP1: DNA Topoisomerase II Binding Protein
UBE4A: Ubiquitination Factor E4A
ZRANB3: Zinc Finger RANBP2-Type Containing 3

## Acknowledgements

We would like to thank Markus Kerr, Nicholas Ariotti, Aaron Smith and Brian Gabrielli for valuable discussion. This work was supported by fellowships and grants from the National Health and Medical Research Council of Australia (to R.G. Parton, (grants APP1140064 and APP1150083 and fellowship APP1156489), R.G. Parton and A.S. Yap, grant number APP1037320, to A.S. Yap, grant number APP1044041, to M.T. Ryan and D.A. Stroud, grant number APP1125390, to D.A. Stroud, grant numbers APP1070916 and APP1140851) as well as by the Australian Research Council Centre of Excellence in Convergent Bio-Nano Science and Technology (R.G. Parton) and the Kids Cancer Project of the Oncology Research Foundation (A.S. Yap). Confocal microscopy was performed at the Australian Cancer Research Foundation (ACRF)/Institute for Molecular Bioscience (IMB) Dynamic Imaging Facility for Cancer Biology, established with funding from the ACRF. The authors acknowledge the use of the Monash Biomedical Proteomics Facility for the provision of instrumentation, training, and technical support. We also thank Beric Henderson (Westmead Institute for Cancer Research, University of Sydney, Australia) for the BRCA1-YFP construct.

## Author Information

KM and RGP conceived the study. KM designed and carried out the experiments, interpreted the data and wrote the manuscript with RGP. DS made the cavin3 KO HeLa cells and performed the LFQ proteomic analysis, YG, ES and MP performed single molecule analysis, MOP preformed preliminary characterization of the knockdown cavin3 A431 and MDA-MB231 cells, KM and GG performed functional experiments. NM performed BRCA1 mRNA analysis, TH cloned BRCA1 into a GFP vector, VT and HP performed preliminary stress experiments, YW performed Western blots, KKK, AY, KA, RD and MR helped design experiments and provided reagents and useful insights. All the authors commented on the manuscript.

## Competing Interests

The authors declare no competing interests.

## Data Availability

All reagents are available from the corresponding author upon request. Proteomics data that supports the findings of this study is presented in Supplementary Table 1. Raw proteomics data will be uploaded to PRIDE upon publication.

## Supplementary Tables

***Supplementary Table 1. Complete LQF proteomics for cavin3 KO cells*.** Complete list of proteins analyzed in cavin3 KO compared to WT HeLa cells (control). Significant (p<0.05) mean log2 transformed SILAC ratios.

***Supplementary Table 2. Pathway analysis for cavin3 KO cells*.** Gene Ontology Biological Process (GOBP) name of both significantly upregulated and downregulated pathways with their corresponding p-values and enrichment scores.

## Supplementary Discussion

In addition to the newly defined roles in homologous recombination and replication stress, cavin3 may play a role in replication fork reversal as several proteins such as HLTF (~2.3 log increase) are significantly upregulated in cavin3-deficient cells. Replication fork reversal is an important protective mechanism that allows forks to reverse their course when DNA lesions are encountered allowing for the resumption of DNA synthesis without chromosome breaks (Neelsen and Lopes, 2015, reviewed in Quintet et al., 2017). Several recent studies by Kolinjivadi et al., 2017; Taglialatela et al., 2017 and Vulganovic et al., 2017 have demonstrated that the translocase activity of ZRANB3 and SMARCAL1 is required for reverse fork formation and is a key function for BRCA1 and BRCA2 in reverse fork protection (Kolinjivadi et al., 2017; Lemacon et al., 2017; Taglialatela et al., 2017 and Vulganovic et al., 2017). ZRANB3 was shown to interact with polyubiquitinated PCNA to promote fork remodeling. Similarly, HLTF was also demonstrated to have DNA translocase activity and to similarly promote PCNA polyubiquitination further implicating HLTF in replication fork reversal (Blastyak et al., 2010). Given the fact that recent studies have suggested that BRCA1 is a crucial regulator of replication fork degradation and that the mechanism leading to fork degradation in these cells remains unclear, further studies are warranted into how different fork remodelers such as ZRANB3 and HLTF work together to mediate replication fork reversal and whether these cells are useful for studying how fork degradation as seen in BRCA1 deficient cells.

Cavin3 KO cells also upregulated proteins that promote the more error-prone Pol/PARP1 mediated Alternate-End Joining (alt-EJ) DNA repair pathway; these included members of the Fanconi anemia (FA) pathway (Fanconi anemia protein component D2 [FANCD2], Fanconi anemia protein component I [FANCI]) as well as protein components of a highly conserved replication stress response pathway such as PARP1, TOPBP1, DROSHA, and RPA2 (**Figure 1**). FANCI was also identified as a potential cavin3 interacting protein in previous studies from our laboratory (McMahon et al., 2019). These findings imply that cavin3 deficient cells would also exhibit increased replication stress and would similarly upregulate the error-prone Pol/PARP1 mediated alt-EJ DNA repair pathway to compensate for defective HR (Ceccaldi et al., 2015; Mateos-Gomez et al., 2015). These findings support further evaluation of replication stress and alternative repair pathways in cavin3 KO cells.

The minichromosome maintenance proteins, MCM2 to MCM7, form a heteromeric DNA helicase required for DNA replication licensing where it primes chromatin for DNA replication (reviewed by Tye et al., 1999; Giaginis et al., 2010). Although DNA helicase activity is required to establish a bidirectional replication fork from each replication origin, a large excess of MCM complexes is accumulated. The role of the additional MCM complexes is not well understood, as most is displaced from the DNA during S-phase, without playing an active role in DNA replication. Recent studies have demonstrated that MCM2-7 expression is downregulated in cells experiencing chronic replication stress and this depends on the tumor suppressor p53 (Bai et al., 2016). A decrease in both HR and NHEJ mediated repair pathways has been demonstrated in cells with a reduction of MCM complex proteins. It was shown that the MCM complex modulates the cellular response to DNA DSBs (Drissi et al., 2018). Indeed, cavin3 KO cells also exhibit downregulation of the MCM proteins, MCM2-7, which may further contribute to the decrease in HR found in these cells. Collectively, these results emphasize the importance of appropriately balancing different repair pathways to maintain global genomic stability and suggest that cavin3 KO cells may be useful for studying these pathways which are particularly relevant in breast and ovarian cancer cells.

---

**Supplementary Figure 1.**
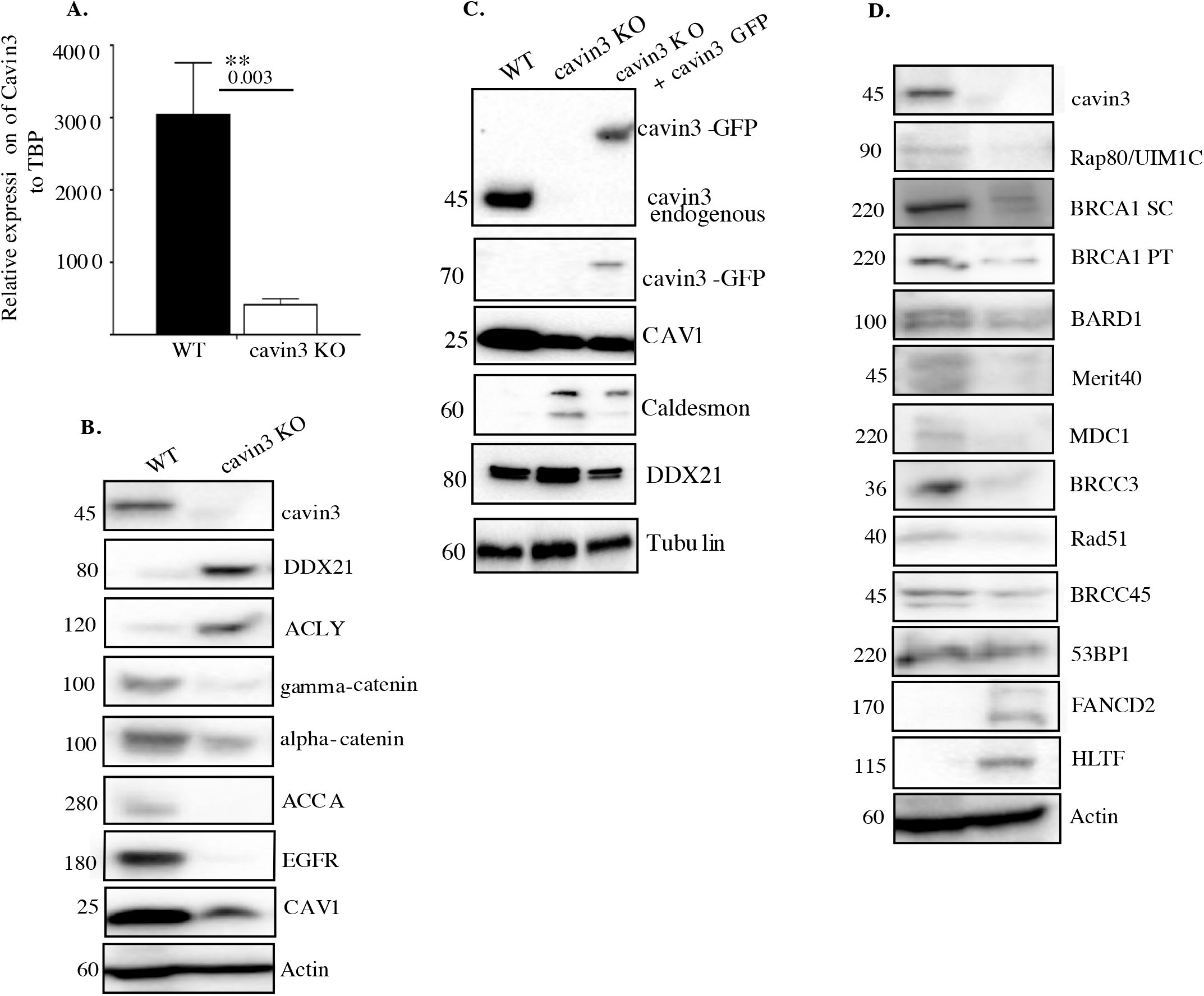
General characterization of cavin3 KO HeLa cells. **(A).** mRNA analysis of cavin3 in WT and cavin3 KO cells from three independent experiments performed in triplicate samples as Mean ± SD using student t-test, ** p<0.01. **(B).** Representative Western blot analysis of equally loaded lysates for cavin3, DDX21, ACLY, gamma-catenin, alpha-catenin, ACCA, EGFR, CAV1 and Actin in WT and cavin3 KO HeLa cell. **(C).** Western blot analysis of equally loaded lysates from WT, cavin3 KO and cavin3 KO transfected with cavin3 for cavin3, GFP, CAV1, Caldesmon, DDX21 and Tubulin as the loading control. Western blots are representative of three independent experiments. **(D).** Representative Western blot analysis of lysates from WT and cavin3 KO cells were Western blotted for cavin3, RAP80/UIM1C, BRCA1 Santa Cruz (SC), BRCA1 Proteintech (PT), BARD1, Merit40, MDC1, BRCC3, Rad51, BRCC45, TP53BP1, FANCD2, HLTF and Actin as the loading control.

**Supplementary Figure 2.**
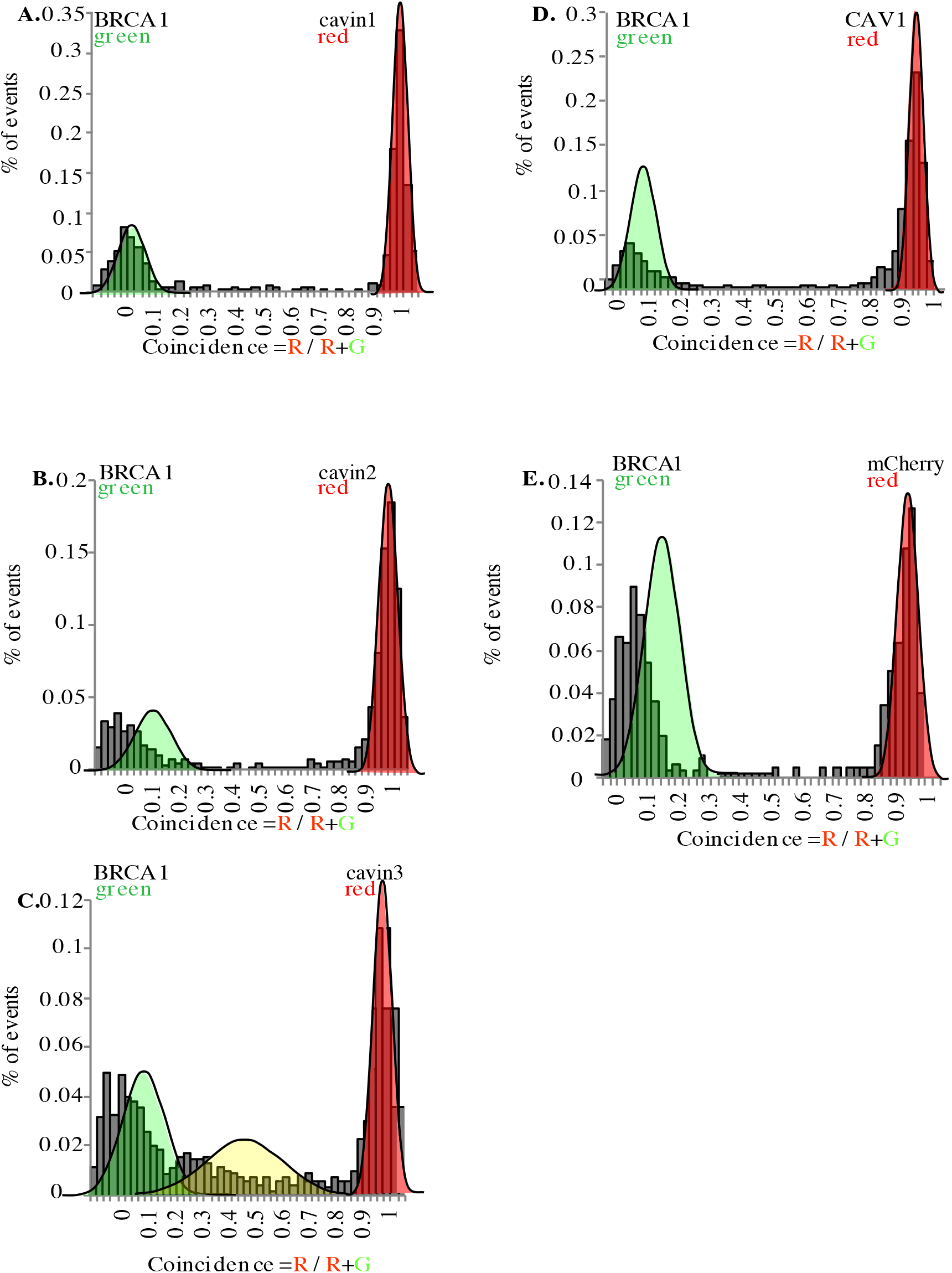
Single molecule analysis in MDA-MB231 cells. Two-color single molecule fluorescence coincidence of BRCA1-GFP with mCherry tagged **(A).** cavin1, **(B).** cavin2, **(C).** cavin3, **(D).** CAV1 and **(E).** mCherry control expressed in MDA-MB231 cells. The green curve represents BRCA1-GFP only events, the red curve represents mCherry only events and the yellow curve represents BRCA1-GFP + Cherry events. More than 1000 events were collected.

**Supplementary Figure 3.**
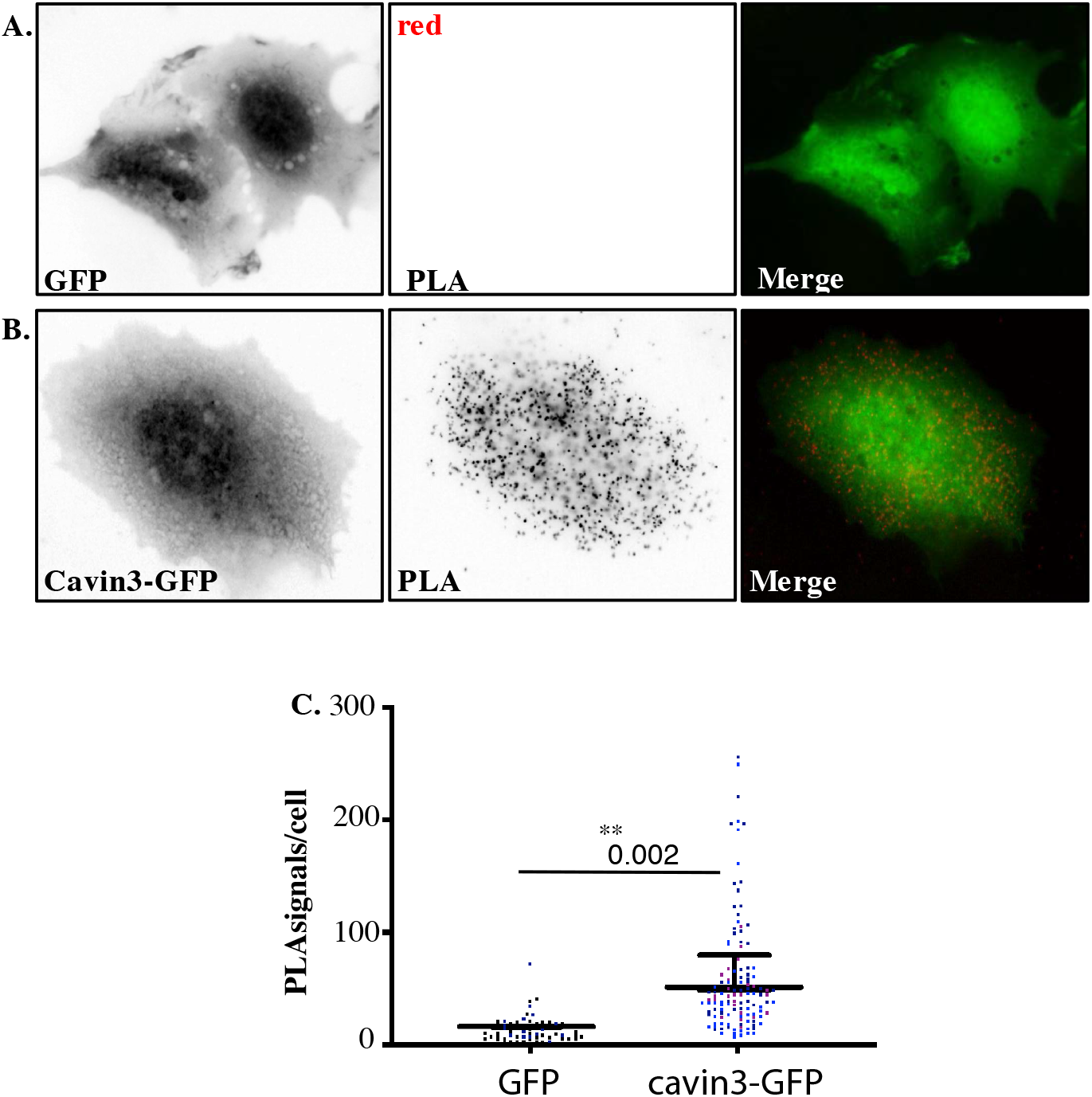
PLA demonstrates Cavin3 and BRCA1 interaction in MCF7 cells. **(A).** Immunofluorescence microscopy in combination with PLA to detect and visualize protein-protein interactions (red dots) within single cells of stably expressing MCF7/GFP and **(B).** MCF7/cavin3-GFP cells using monoclonal BRCA1 (MS 110) and polyclonal GFP antibodies. DNA was counterstained with DAPI (blue). Scale bars represent 10 μm. **(C)**. Number of red dots/PLA signals in 40-50 cells for each MCF7/GFP expressing cell line quantified from 3 independent experiments using a nested ANOVA. Each biological replicate is color coded with the Mean ± SEM presented as a black bar. ** p<0.01.

**Supplementary Figure 4.**
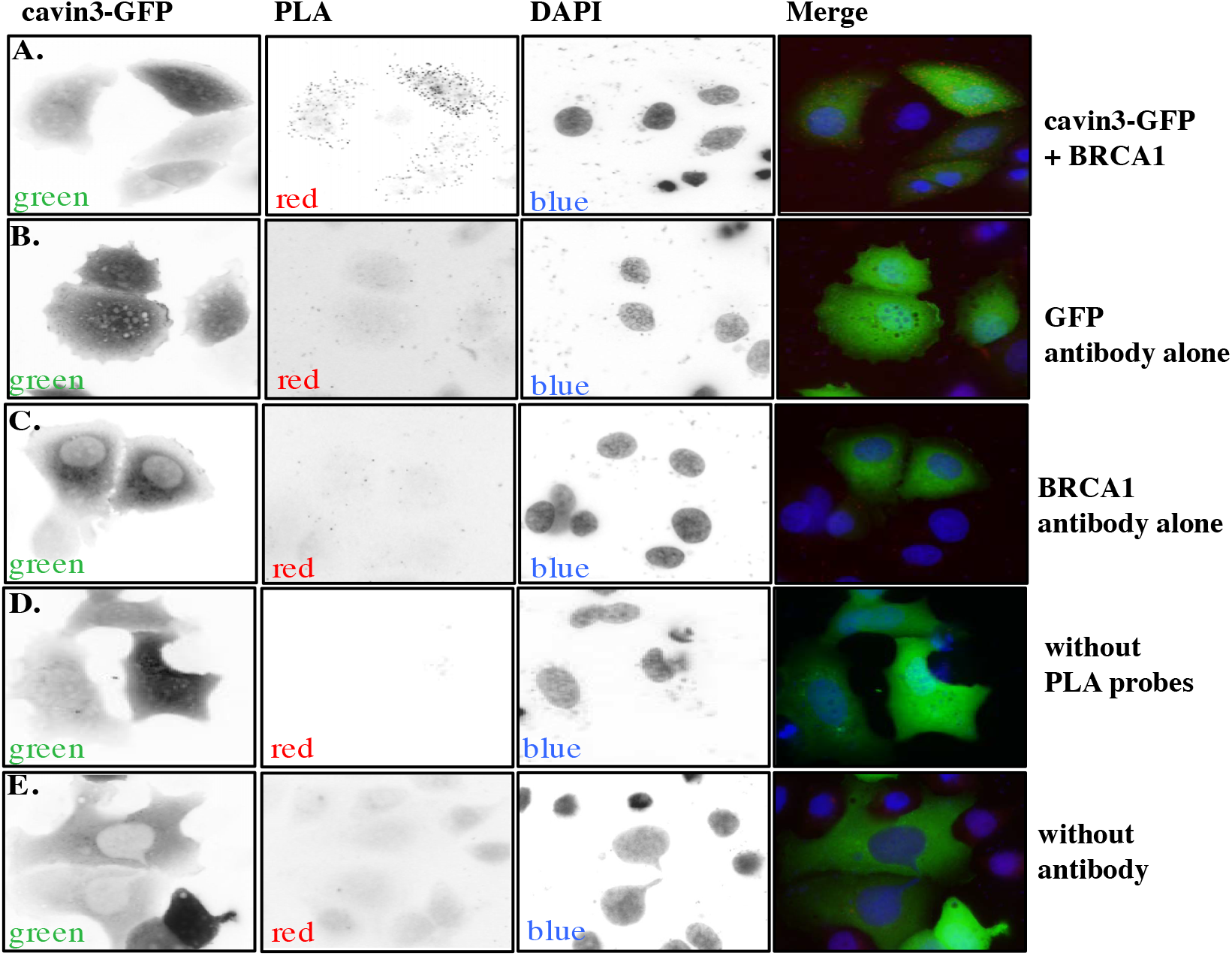
PLA controls. **(A).** Fluorescence microscopy analysis of PLA signals generated in MCF7 cells transfected with cavin3-GFP using mouse GFP and rabbit BRCA1 antibodies from Figure 3D, **(B).** GFP antibody alone, **(C).** BRCA1 antibody alone, **(D).** the absence of PLA probes or **(E).** primary antibody controls. Representative images are from at least two independent experiments as shown.

**Supplementary Figure 6.**
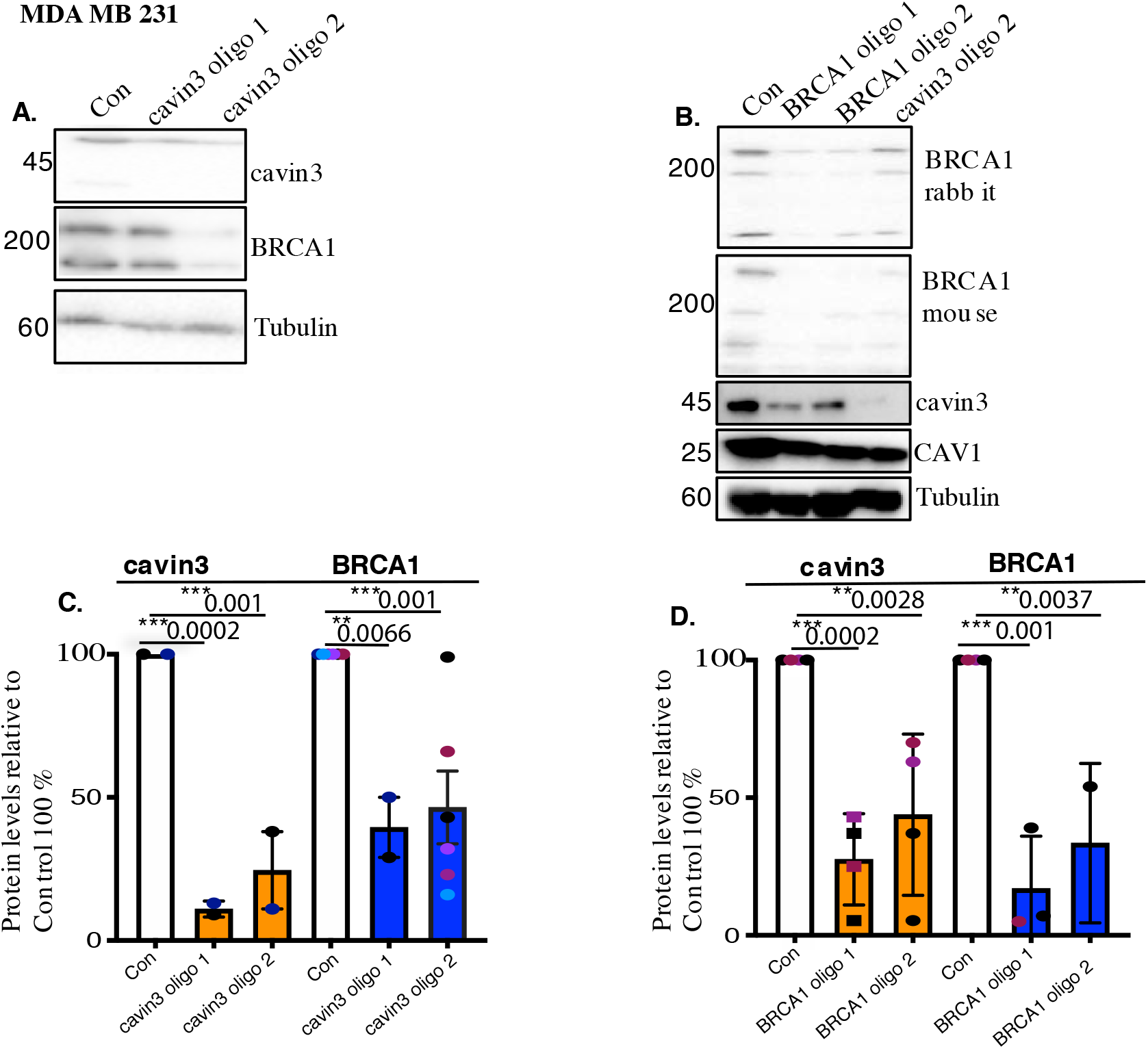
Validation of loss of Cavin3 and BRCA1 in MDA-MB231 cells. **(A).** Representative Western blot analysis of cavin3, BRCA1, CAV1 and Tubulin as the loading control in MDA-MB231 cells following treatment with two siRNAs targeting cavin3 oligo 1 and 2 (or control). **(B)**. Relative protein expression of cavin3 and BRCA1 in MDA-MB 231 cells transfected with cavin3 oligo 1 and cavin3 oligo 2 compared to control treated cells from three independent experiments using a Oneway ANOVA with Bonferroni multiple comparison tests. Each biological replicate is color coded. **(C).** Western blot analysis of BRCA1, cavin3 and Tubulin as the loading control in MDA-MB 231 cells following treatment with two siRNAs targeting BRCA1 oligo 1 and 2 (or control). **(D)**. Relative protein expression of cavin3 and BRCA1 in MDA-MB 231 cells transfected with cavin3 oligo 1 and cavin3 oligo 2 compared to control treated cells from three independent experiments using a one-way ANOVA with Bonferroni multiple comparison test. Each biological replicate is color coded. ** p<0.01, ***p<0.001.

**Supplementary Figure 5.**
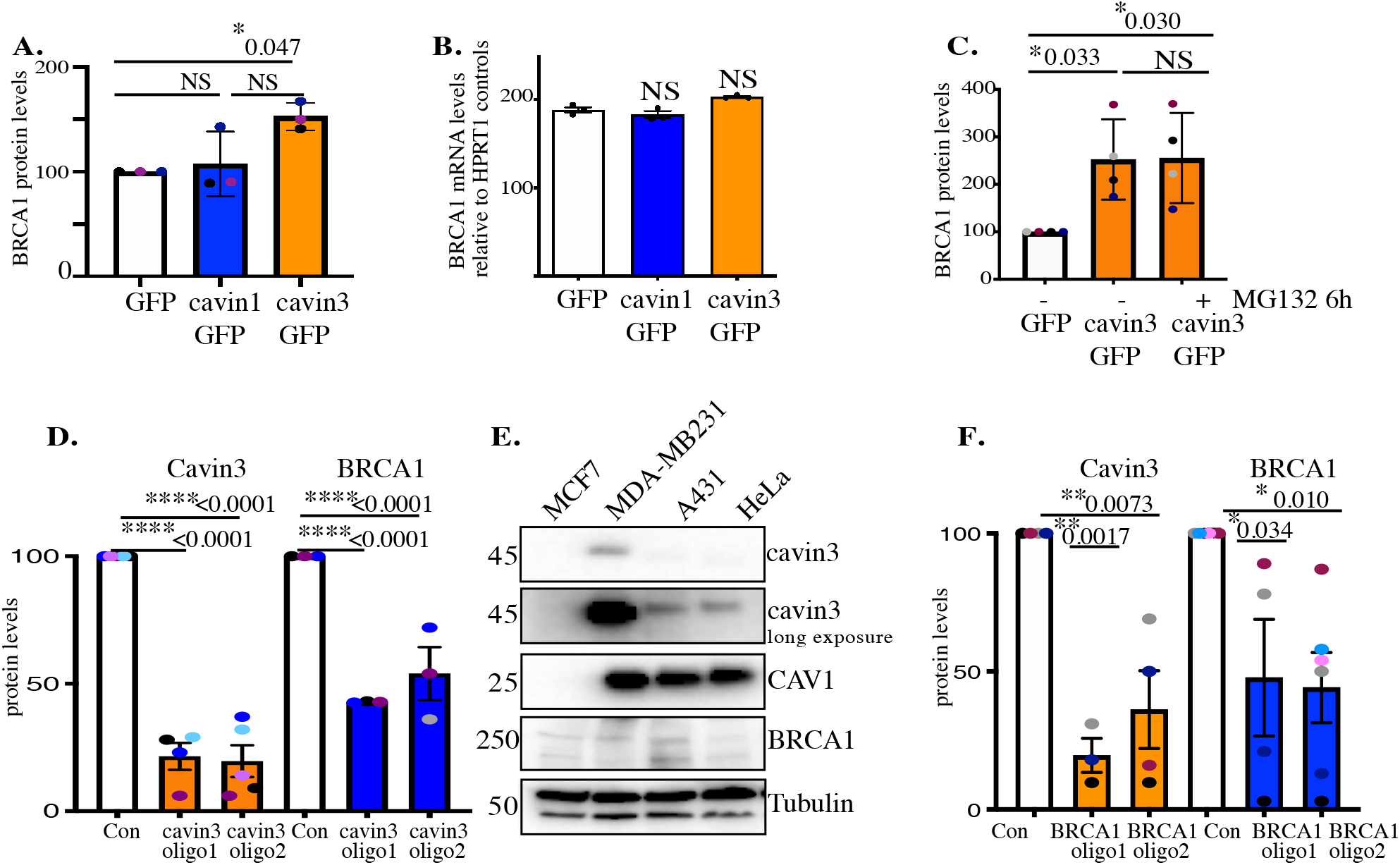
Reciprocal regulation of BRCA1 and cavin3 protein levels. **(A).** Relative protein expression of BRCA1 in MCF7 cells expressing GFP (white), cavin1-GFP (blue) and cavin3-GFP (orange) using a one-way ANOVA and Bonferroni’s multiple comparisons test from three independent experiments. **(B).** BRCA1 gene expression was analysed using the Taqman Gene Expression assay as described in Materials and Methods in MCF7 cells. Results are expressed as Mean ± SD using a one-way ANOVA and Bonferroni’s multiple comparisons test from three independent experiments. NS, not significant. **(C).** Relative protein expression of BRCA1 in MCF7 cells expressing GFP or cavin3-GFP untreated (-) or treated with MG132 (+) using a one-way ANOVA and Bonferroni’s multiple comparisons test from three independent experiments. **(D).** Relative protein levels of cavin3 and BRCA1 in cells treated with control (Con) siRNAs or siRNAs specific to cavin3 (oligo1 and oligo2) in A431 cells using a one-way ANOVA and Bonferroni’s multiple comparisons test from three-four independent experiments. **(E).** Representative Western blots of MCF7, MDA-MB231, A431 and HeLa cells were Western blotted for protein expression of cavin3, CAV1, BRCA1 and Tubulin as the loading control **(F).** Relative protein levels of cavin3 and BRCA1 in cells treated with control (Con) siRNAs or siRNAs specific to BRCA1 (oligo1 and oligo2) in A431 cells using a one-way ANOVA and Bonferroni’s multiple comparisons test from three-four independent experiments. For each experiment, each biological replicate was color coded. NS – not significant, * p<0.05, ** p<0.01, ***p<0.001, **** p<0.0001

**Supplementary Figure 7.**
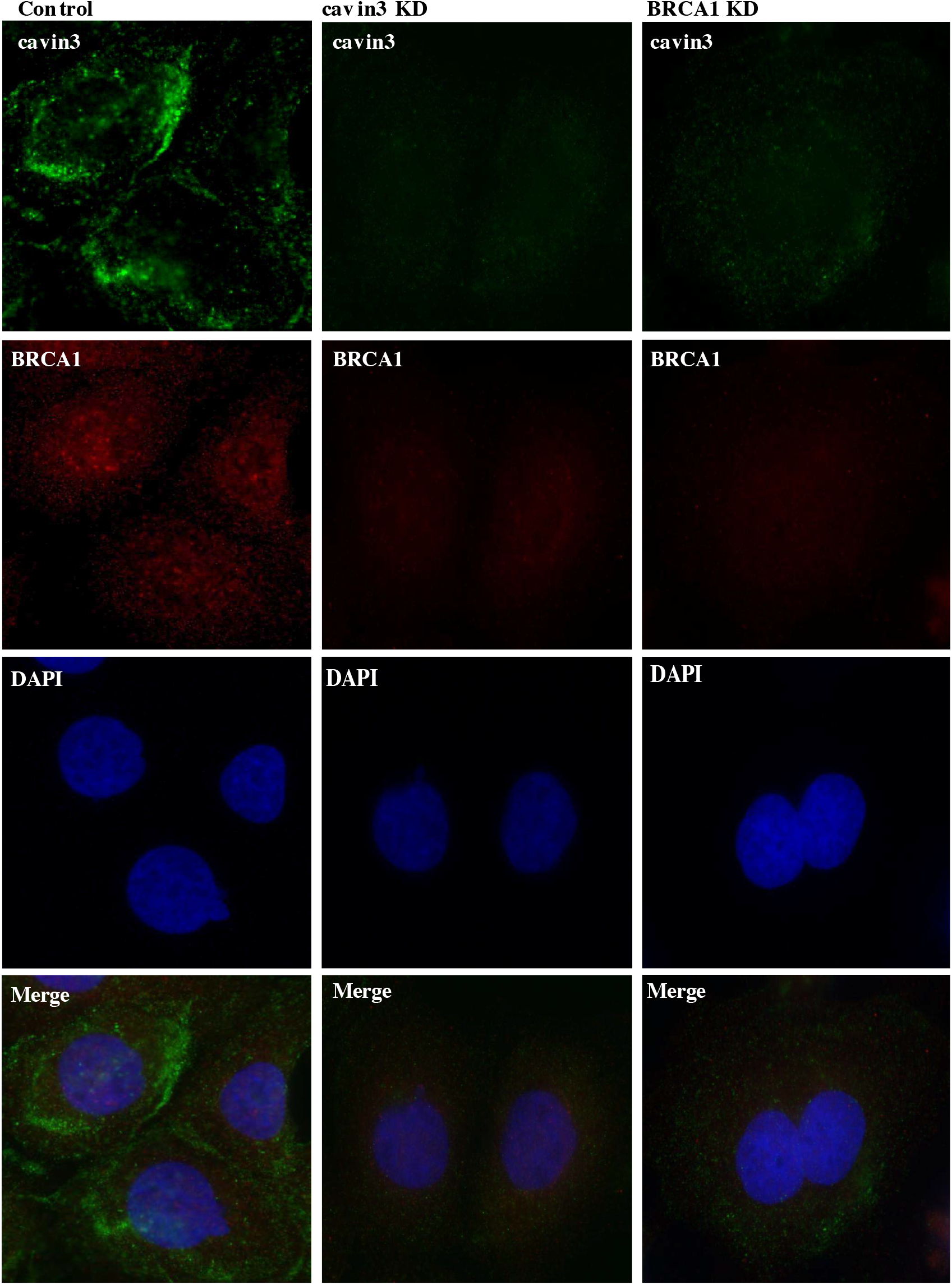
Reciprocal loss of BRCA1 and cavin3 in A431 cells. Representative immunofluorescence images of Control knockdown, cavin3 knockdown and BRCA1 knockdown cells for cavin3 (green), BRCA1(red) and DAPI (nuclei). Images are representative of three independent experiments.

**Supplementary Figure 8.**
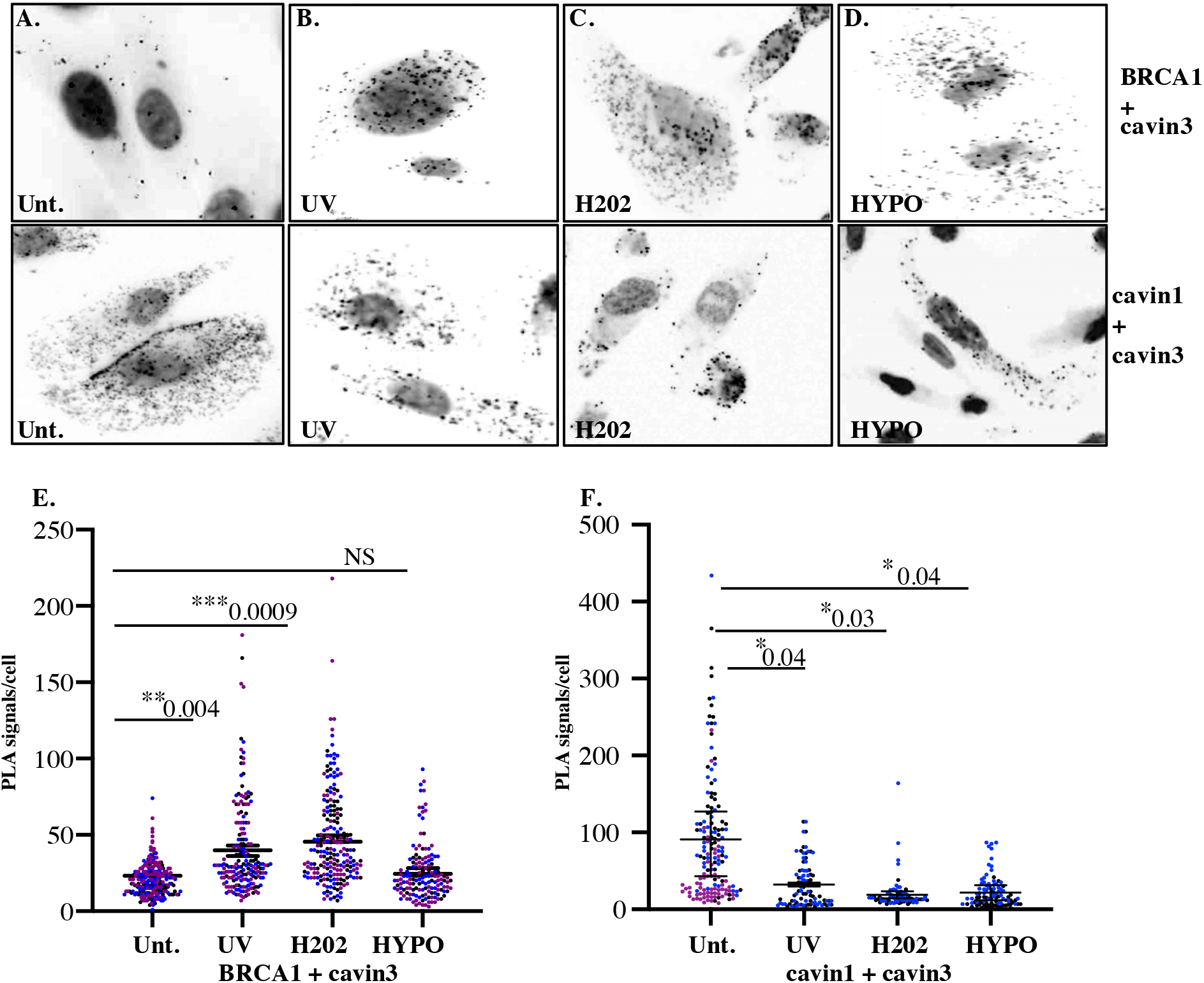
Close association of Cavin3 and BRCA1 in MDA-MB231 cells after stress treatment. **(A).** Immunofluorescence microscopy in combination with PLA visualization of endogenous protein–protein interactions (red dots) within MDA-MB231 cells in untreated (Unt.) cells, **(B).** UV treatment (2 min) and a 30 min chase time, **(C).** 200 μM H_2_0_2_ for 30 min and **(D).** Hypo-osmotic treatment (90% H_2_0 in DMEM) for 10 min. **(E).** Number of red dots/PLA signals in 40-50 cells for cavin-BRCA1 was quantified from 3 independent experiments and is presented as Mean ± SEM using a nested ANOVA. Each biological replicate is color coded with the mean presented as a black bar. **(F).** Number of red dots/PLA signals in 40-50 cells for cavin3-cavin1 was quantified from 3 independent experiments and is presented as Mean ± SEM using a nested ANOVA. Each biological replicate is color coded with the mean presented as a black bar. * p<0.05, ** p<0.01, ***p<0.001.

**Supplementary Figure 9.**
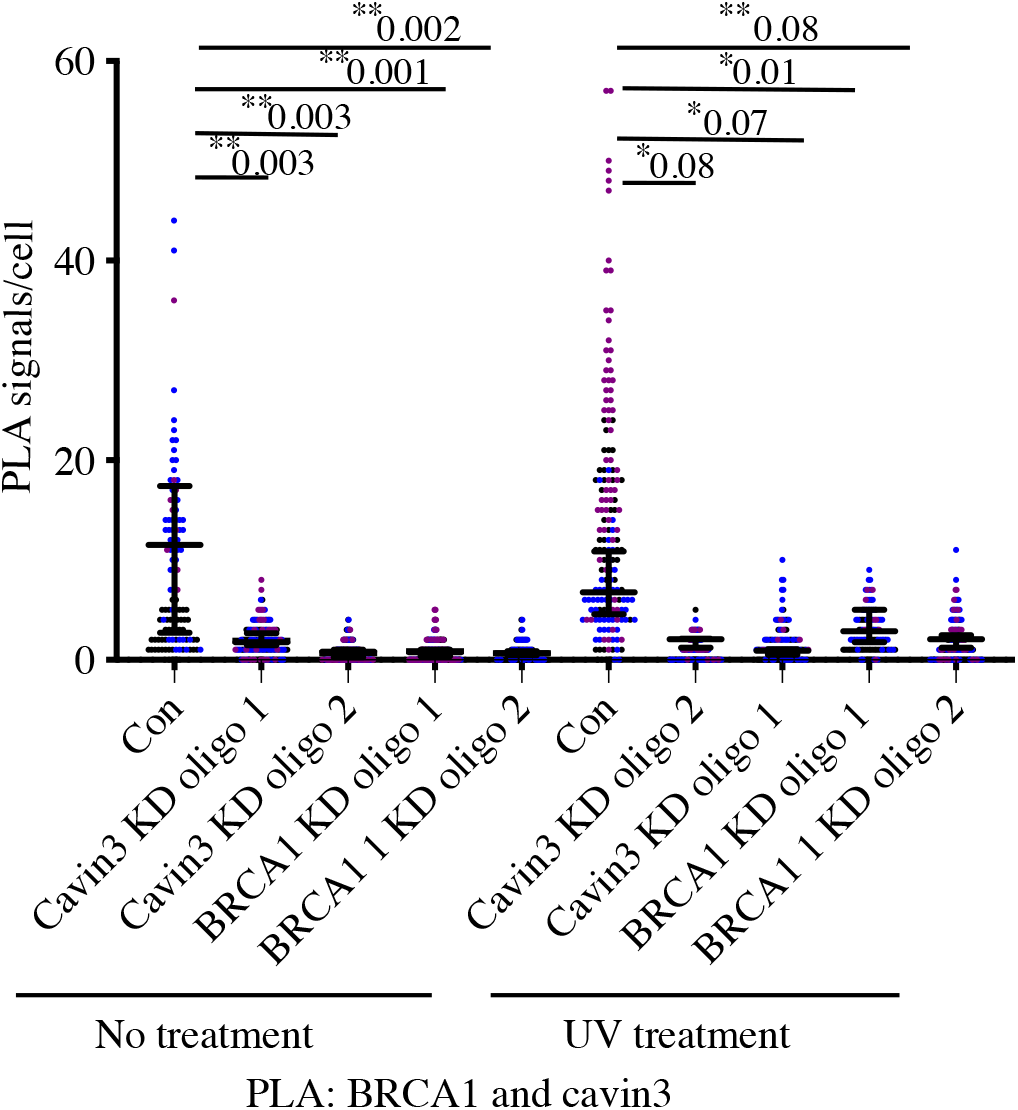
PLA controls for cavin3 and BRCA1 PLA antibodies in A431 cells. A431 cells treated with control (Con) or siRNAs specific to cavin3 or BRCA1 (oligo1 and 2). Cells were left untreated or subjected to UV treatment. Cells were subjected to immunofluorescence microscopy in combination with PLA (red dots) using monoclonal BRCA1 and polyclonal cavin3 antibodies. The number of red dots/PLA signals in 40-50 cells was quantified from 3 independent experiments using a nested ANOVA. Each biological replicate is color coded with the Mean ± SEM presented as a black bar. *p<0.05, **p<0.01.

**Supplementary Figure 10.**
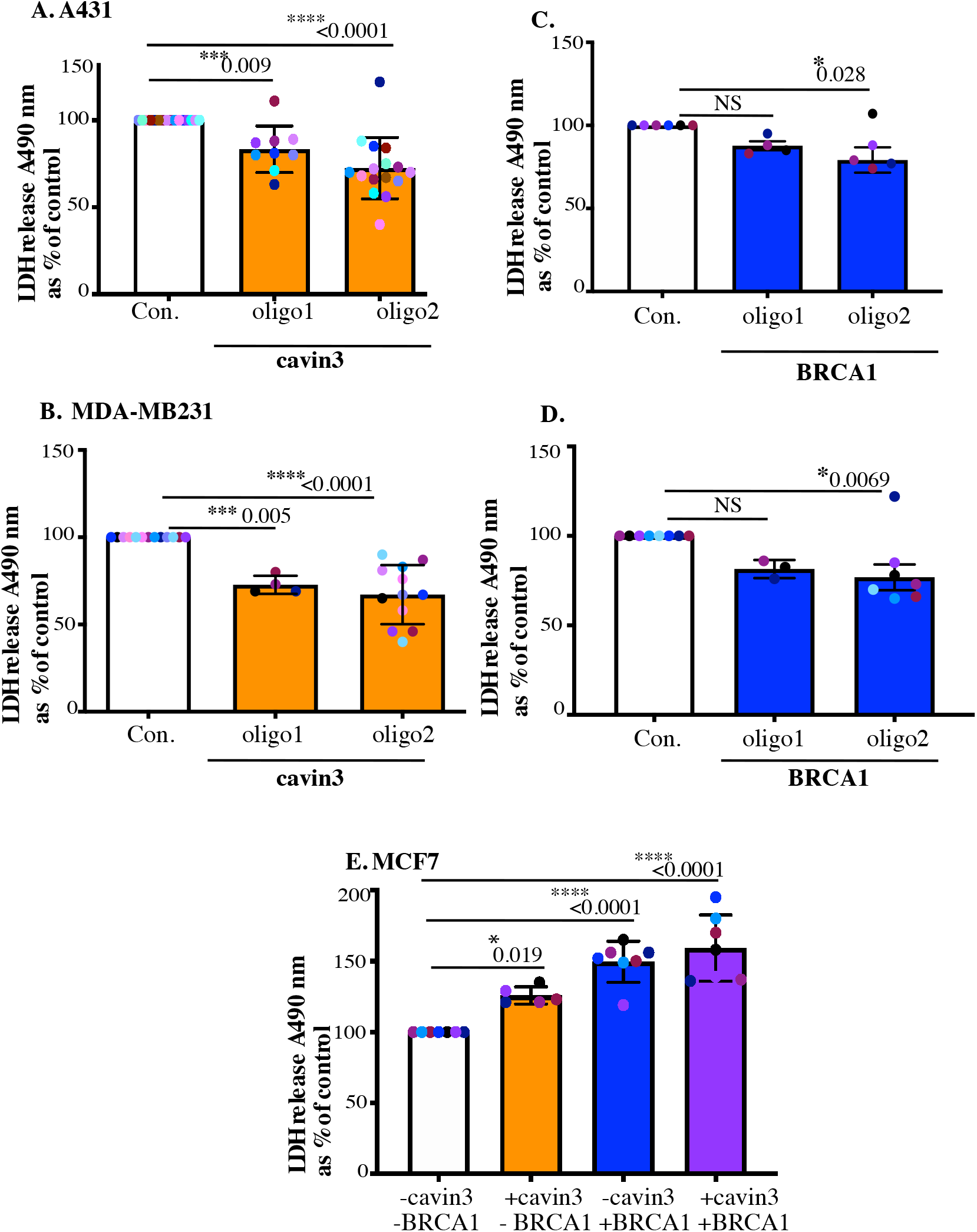
Validation of LDH release in MCF-7, A431, and MDA-MB231 cells. **(A).** Equal numbers of A431 cells treated with control siRNA, or cavin3 specific siRNAs were subjected to UV treatment and a recovery time of 6 hours. **(B).** Equal numbers of A431 cells treated with control siRNAs or BRCA1 specific siRNAs were subjected to UV treatment (2 min) and a recovery time of 6 hours. **(C).** Equal numbers of MDA-MB231 cells treated with control siRNAs or cavin3 specific siRNAs were subjected to UV treatment and a recovery time of 6 hours. **(D).** Equal numbers of MDA-MB231 cells treated with control siRNAs or BRCA1 specific siRNAs were subjected to UV treatment and a recovery time of 6 hours. LDH release was measured from the cell supernatant and was calculated relative to control siRNA UV treated cells as Mean ± SD using a one-way ANOVA and Bonferroni’s multiple comparisons test. **(E).** MCF7 cells depleted of BRCA1 (-cavin3, -BRCA1), depleted of BRCA1 and transfected with cavin3-GFP (-BRCA1, + cavin3), left untreated (-cavin3, + BRCA1) or transfected with cavin3-GFP (+ BRCA1, + cavin3). All cells were subjected to UV treatment and a 6 hour recovery time. LDH release was measured from the cell supernatant and was calculated relative to control MCF7 cells lacking both BRCA1 and cavin3 as Mean ± SD using a one-way ANOVA and Bonferroni’s multiple comparisons test. Each biological replicate is color coded. NS – not significant, * p<0.05, ***p<0.001, **** p<0.0001.

**Supplementary Figure 11.**
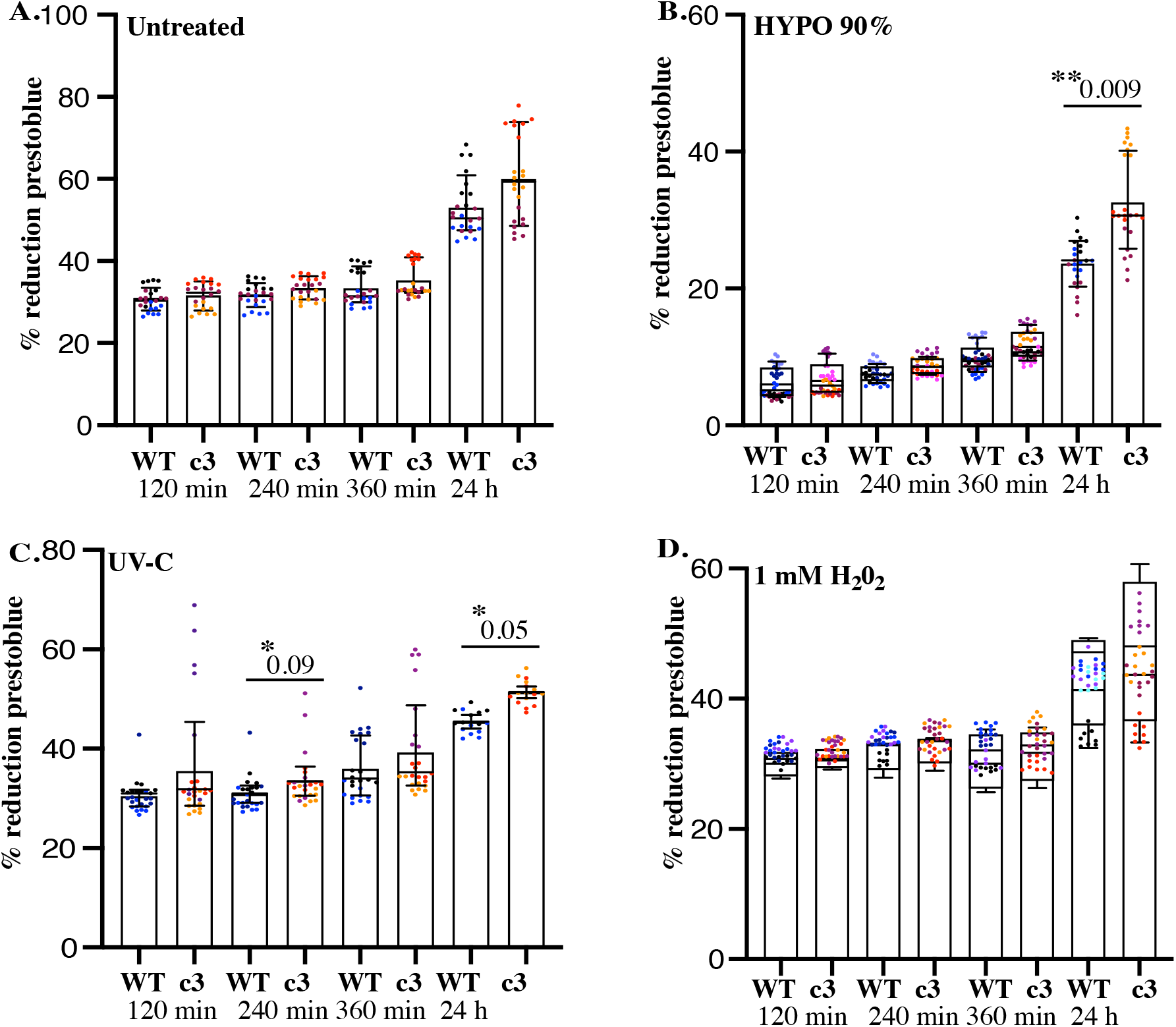
Cavin3 KO cells exhibit resistance to stressors that allow BRCA1 interaction. Equal numbers of WT and cavin3 KO cells were either left **(A).** untreated, **(B).** treated with 90% hypo-osmotic medium, **(C).** UV-C 2 min or **(D).** 1 mM H_2_0_2_ (oxidative stress). Presto blue reagent was added to plates immediately and were read at 570 and 600 nm at 120 min, 240 min, 360 min and 24 hours. The % reduction prestoblue was calculated from eight wells/replicate experiment and is presented as the Mean ± SEM using a nested ANOVA for each time point from three-four independent experiments. Each biological replicate is color coded. * p<0.05, **p<0.01.

**Supplementary Figure 12.**
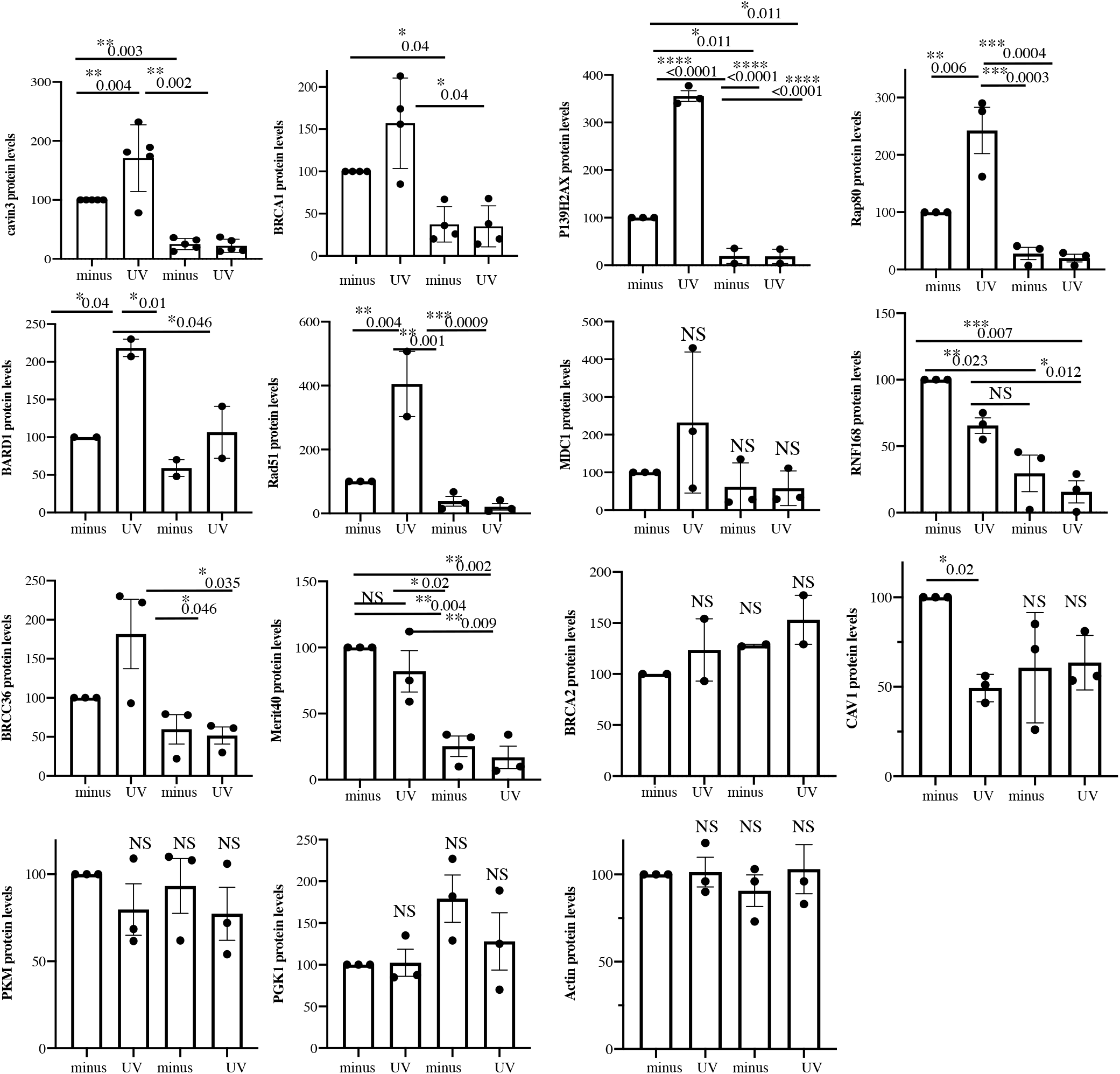
Quantitation of BRCA1-A-complex proteins in WT and cavin3 KO cells. Densitometry analysis was performed of the protein levels of cavin3, BRCA1, P139 γH2AX, RAP80, BARD1, RAD51, MDC1, RNF168, BRCC36, Merit40, BRCA2, CAV1, PKM, PGK1 and Actin in Figure 8A and 8C in WT and cavin3 KO cells subjected to UV treatment and a 4 hour chase from two-three independent experiments presented as Mean ± SD using a one way ANOVA and Bonferroni’s multiple comparisons test. NS, not significant, * p<0.05, ** p<0.01, ***p<0.001, **** p<0.0001.

## Notes

### Competing Interest Statement

The authors have declared no competing interest.

